# Cytoklepty in the plankton: a host strategy to optimize the bioenergetic machinery of endosymbiotic algae

**DOI:** 10.1101/2020.12.08.416644

**Authors:** Uwizeye Clarisse, Mars Brisbin Margaret, Gallet Benoit, Chevalier Fabien, LeKieffre Charlotte, Schieber L. Nicole, Falconet Denis, Wangpraseurt Daniel, Schertel Lukas, Stryhanyuk Hryhoriy, Musat Niculina, Mitarai Satoshi, Schwab Yannick, Finazzi Giovanni, Decelle Johan

## Abstract

Endosymbioses have shaped the evolutionary trajectory of life and remain widespread and ecologically important. Investigating modern oceanic photosymbioses can illuminate how algal endosymbionts are energetically exploited by their heterotrophic hosts, and inform on putative initial steps of plastid acquisition in eukaryotes. By combining 3D subcellular imaging with photophysiology, carbon flux imaging and transcriptomics, we show that cell division of algal endosymbionts (*Phaeocystis*) is blocked within hosts (Acantharia), and that their cellular architecture and bioenergetic machinery are radically altered. Transcriptional evidence indicates that a nutrient-independent mechanism prevents symbiont cell division and decouples nuclear and plastid division. As endosymbiont plastids proliferate, the volume of the photosynthetic machinery volume increases 100-fold in correlation with expansion of a reticular mitochondrial network in close proximity to plastids. Photosynthetic efficiency tends to increase with cell size and photon propagation modeling indicates that the networked mitochondrial architecture enhances light capture. This is accompanied by 150-fold higher carbon uptake and upregulation of genes involved in photosynthesis and carbon fixation, which, in conjunction with a ca.15-fold size increase of pyrenoids demonstrates enhanced primary production in symbiosis. NanoSIMS analysis revealed major carbon allocation to plastids and transfer to the host cell. Invagination of the symbiosome into endosymbionts to optimize metabolic exchanges is strong evidence that the algal metamorphosis is irreversible. Hosts therefore trigger and unambiguously benefit from major bioenergetic remodeling of symbiotic microalgae with important consequences for the oceanic carbon cycle. Unlike other photosymbioses, this interaction represents a so-called cytoklepty, which is a putative initial step towards plastid acquisition.

## Introduction

Living with intracellular microalgae (photosymbiosis) is a globally widespread and ecologically important lifestyle. In marine benthic ecosystems, a wide diversity of animals (e.g. cnidarians, molluscs) host photosynthesizing microalgae in their tissues, sustaining coral reef habitats and their associated biodiversity worldwide. In the oceanic plankton, various unicellular heterotrophic eukaryotes (e.g. radiolarians, foraminiferans) also establish symbioses with intracellular microalgae (1, 2). These ubiquitous organisms contribute significantly to primary production (3) and carbon sequestration from surface waters to the deep sea (4–7). Photosymbioses are generally recognized as mutualistic partnerships with hosts providing nutrient-rich and protective microhabitats to algal symbionts, which in return provide energy-rich compounds produced via photosynthesis (e.g. sugars, lipids; (8)). This metabolic crosstalk between a heterotroph and an autotroph provides a competitive advantage in nutrient-poor habitats, such as the open ocean. However, it is difficult to demonstrate whether photosymbioses confer evolutionary advantages on algal symbionts, particularly in marine planktonic symbioses, which cannot be cultured in laboratory conditions and where the costs and benefits for each partner are not clearly defined (9). Therefore, whether photosymbioses are true mutualistic partnerships has long been debated (10).

In the evolutionary history of eukaryotes, photosymbiosis is recognized as a preliminary step leading to plastid acquisition and spread across photosynthetic lineages (11, 12). Interactions between host and endosymbiont evolved into inverted parasitism whereby the host exploited a prokaryotic (primary endosymbiosis) or plastid-bearing eukaryotic symbiont (secondary or tertiary endosymbiosis), leading to gradual cellular and genomic reduction of the symbiont and ultimately plastid acquisition by the host (13–15). However, the underlying cellular mechanisms that allow a host cell to take control and manipulate the bioenergetics of intracellular microalgae remain unknown. Several relevant models have been studied, but reduction of the microalgal cell and horizontal gene transfer to the host has already occurred in each of them (e.g. kleptoplastidy in dinoflagellates (16), *Paulinella chromatophora* (17)). Studying contemporary unicellular photosymbioses involving intact microalgae has the potential not only to lead to a better understanding of their ecological success in the ocean but also to provide evolutionary insights into the putative first steps of plastid acquisition and underlying host control.

The cosmopolitan planktonic photosymbiosis between acantharian hosts and intact microalgae (*Phaeocystis spp.,* Haptophyta) is characterized by a morphological transformation of symbionts, wherein algal cell volume increases dramatically and there is a multiplication of enlarged plastids (18, 19). Free-living *Phaeocystis* cells can reach high population densities in many oceanic regions (20, 21), thus confounding any evolutionary advantages symbiosis may confer on this microalga. Conversely, the ecological success of free-living *Phaeocystis* benefits hosts since it increases opportunities to capture new symbionts throughout the life cycle (22). Hence, Acantharia-*Phaeocystis* photosymbioses are potentially exploitive symbioses with intact symbiont cells and represent an ideal system to investigate this oceanic interaction and bring new insights into the early transitional stages of more permanent algal endosymbioses.

To elucidate mechanisms involved in host exploitation of algal cells and the putative initial steps of plastid acquisition, the structural, physiological, and genetic changes of symbiotic microalgae need to be explored at the subcellular and molecular level. Here, the subcellular architecture of *Phaeocystis* cells outside (i.e. free-living in culture) and inside host cells was reconstructed in 3D to quantify structural changes of energy-producing organelles (plastid, mitochondria) and their interactions. In parallel, single-cell transcriptomics compared *Phaeocystis* gene expression in free-living and symbiotic stages. This combination of quantitative subcellular imaging and transcriptomics showed that the cell cycle of endosymbiotic microalgae is halted and the bioenergetic machinery is drastically enhanced. We observed a proliferation of plastids with enlarged pyrenoids and correlated extension of reticulated mitochondria in symbionts, accompanied by the upregulation of genes involved in photosynthesis and central carbon pathways. Photophysiology, photon propagation modeling and carbon flux imaging further demonstrated that algal energy production is significantly enhanced, leading to increased production of organic carbon, which is allocated to plastids and transferred to the host cell. This study deciphers mechanisms on how a microalgal cell is morphologically, metabolically and transcriptionally modified and ultimately exploited by a heterotrophic host cell in the oceanic plankton (i.e. cytoklepty).

## Results and Discussion

### *Phaeocystis* cell division is inhibited in symbiosis

Using 3D electron microscopy (FIB-SEM: Focus Ion Beam - Scanning Electron Microscopy), we reconstructed and performed morphometric analyses of the subcellular architecture of the microalga *Phaeocystis* in the free-living (n = 20 cells) and endosymbiotic stage within two distinct acantharian hosts (n = 7 cells) (Fig. 1, Dataset S1). Alongside the disappearance of flagella and external scales, the total volume of symbiotic *Phaeocystis* cells increased 6- to 78-fold compared to free-living cells and accommodated more organelles and larger vacuoles (Figs. 1 and 2). Such a dramatic increase in cell size indicates that *Phaeocystis* cell-division could be blocked while living symbiotically (23, 24). Consistent with this hypothesis, single-cell transcriptome analysis on twelve distinct hosts revealed that key *Phaeocystis* genes involved in DNA replication and progression through cell cycle stages (G1, S, G2, M) were downregulated in symbiosis (Fig. 2 and Fig. S1), including genes for DNA polymerase complexes, cyclins, cyclin-dependent kinases, and the anaphase-promoting complex. Additionally, nuclear volume increased in symbiotic cells up to 32-fold (Fig. 2, Dataset S1), although its relative occupancy decreased from 9.3 ± 1.6% in free-living cells to 4.7 ± 2% in symbiosis (Fig. 1D). Taking advantage of the varying electron densities of compartments within the nucleus, we separately quantified the volumes of the nucleolus (site of ribosome genesis) (25) and, eu- and hetero-chromatin (sites of DNA replication and gene expression) (26). The volume of these three nuclear compartments concomitantly increased in symbiosis (20-fold for the nucleolus, 18-fold for the heterochromatin, 23-fold for euchromatin) and their volume ratios remained relatively stable (Fig. S1, Dataset S1). While there is no reorganization of chromatin in symbiosis, the overall increase of its volume suggests that symbionts accumulate DNA. DNA synthesis occurs in symbiosis, but the final steps of mitosis and cytokinesis are prevented. Increased cell and nuclear volume alongside downregulation of DNA replication and cell-cycle pathways compile strong evidence for inhibited cell division in symbiotic *Phaeocystis*.

**Figure 1.**
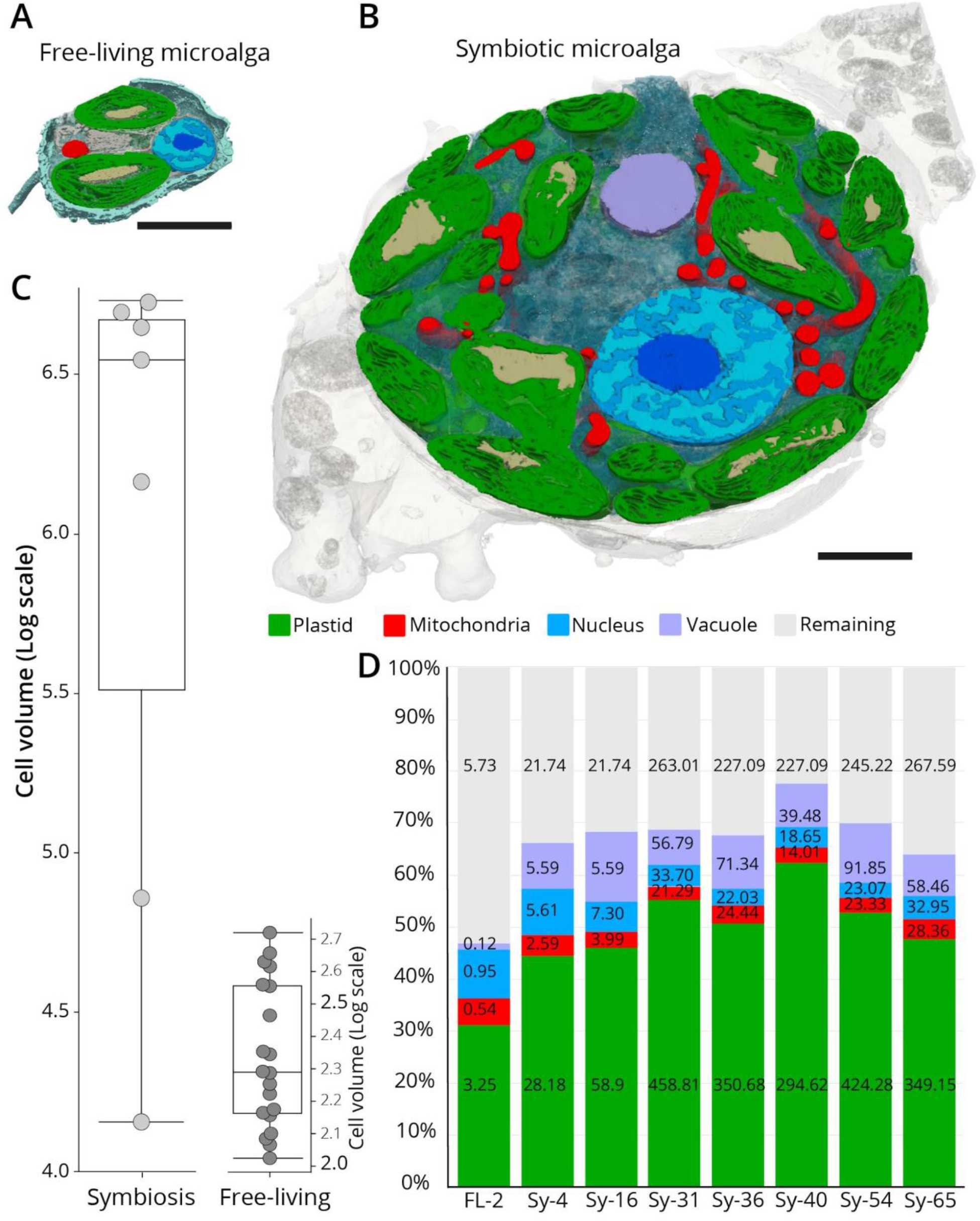
Morphological transformation of the microalga *Phaeocystis cordata* in symbiosis unveiled by FIB-SEM. **A–B)** Sections of the 3D reconstruction of the free-living (A) and symbiotic (B) *Phaeocystis* cells as revealed by FIB-SEM (Focused Ion Beam-Scanning Electron Microscopy), showing multiplication of plastids (green) with immersed pyrenoids (light brown), extension of the mitochondria (red), nuclear compartments (nucleolus in dark blue, heterochromatin in blue, and euchromatin in light blue), and vacuoles (purple). Scale bar: 2 μm. **C)** Box plots showing the increase of the cell volume (log scale, μm^3^) in symbiotic *Phaeocystis* (left) compared to the free-living cells (right). **D)** Relative volume occupancy of different organelles and cellular compartments (plastid, mitochondria, nucleus, vacuole) as % occupancy in the cell (organelle volume/cell volume ratio) in free-living (FL-2 plastids) and seven different symbiotic microalgal cells (Sy) having 2, 4, 16, 31, 36, 40, 54, 65 plastids. The volumes (μm^3^) of organelles and cellular compartments are given within respective bar segments, and in Dataset S1

**Figure 2.**
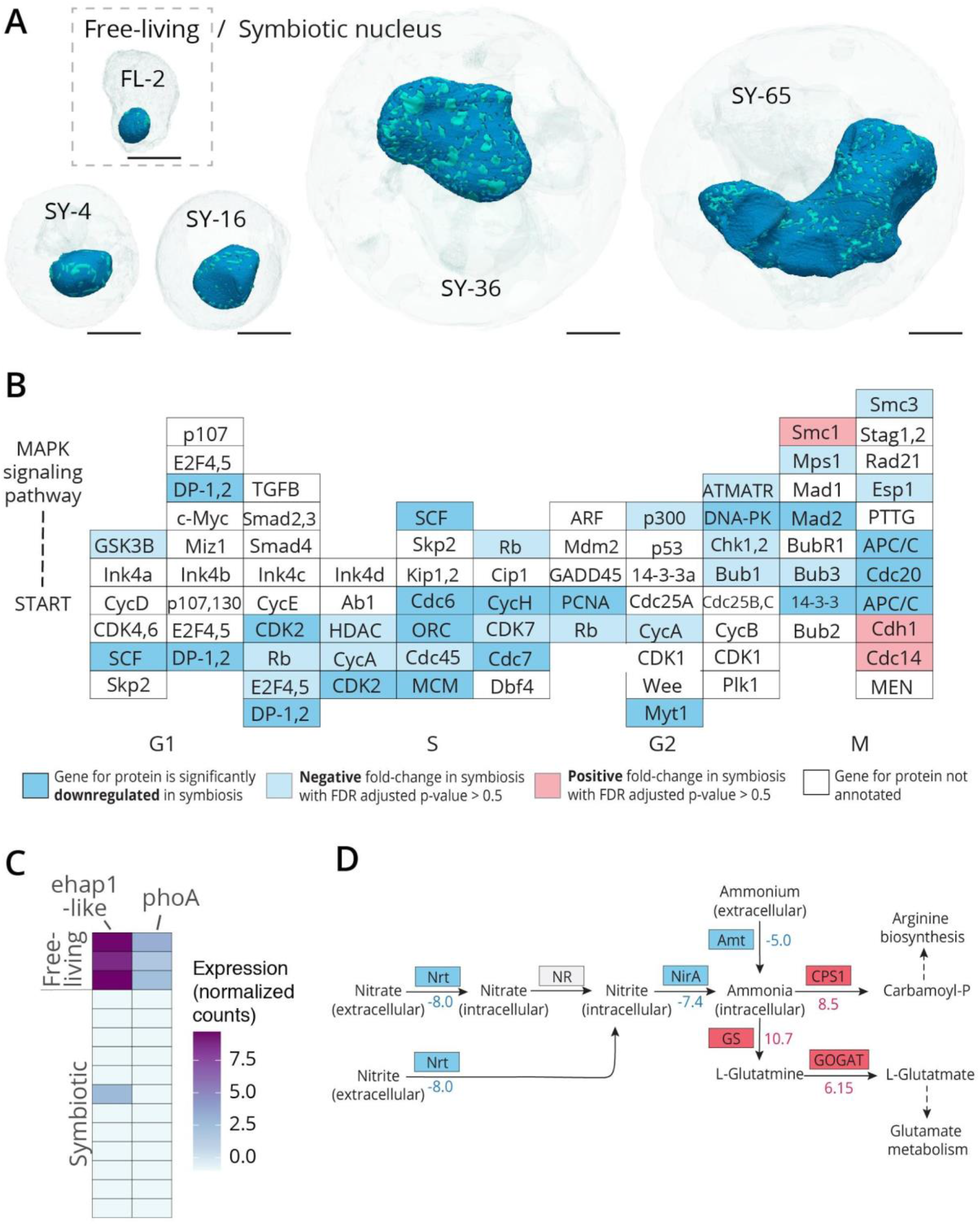
Increased nuclear volume and inhibited cell cycle in symbiotic *Phaeocystis* despite lack of evidence for nutrient limitation. **A)** 3D reconstruction of algal cells (transparent) and their nucleus (blue) after FIB-SEM imaging, showing increased nuclear volume in symbiotic algal cells with 4, 16, 36, 65 plastids from left to right, compared to the free-living stage (top left) (See also Fig. S1). Scale bar: 2 μm **B)** Differential expression of *Phaeocystis* genes in the KEGG cell cycle reference pathway (ko04110). Blocks representing protein and enzymes are colored according to differential gene expression results. Solid blue indicates genes that were significantly downregulated in symbiosis (padj < 0.05, log2FC < −1); transparent blue indicates genes that had a negative fold-change in symbiosis but the difference was not statistically significant; transparent red indicates genes that had a positive, but statistically insignificant, fold-change in symbiosis; white indicates proteins without annotated genes. **C)** Heatmap of alkaline phosphatase gene expression (*phoA* and *ehap1*-like) as log normalized counts in free-living *Phaeocystis cordata* cultures (n = 3) and in individual hosts (n = 12). Alkaline phosphatase expression is a marker for phosphorus limitation in microalgae and these genes were mostly not expressed in symbiosis. **D)** Differential expression of genes in the Nitrogen metabolism KEGG reference pathway (ko00910). Inorganic nitrogen transporters, especially those encoded by *Nrt*, are marker genes for nitrogen limitation in microalgae. Genes that were significantly downregulated in symbiosis are colored blue and those that were significantly upregulated are red. The log2 fold-change values for differentially expressed genes are indicated next to or below the gene name.

Preventing symbiont cell division is a host strategy for managing symbiont populations and limiting symbiont overgrowth (27). Hosts can regulate symbiont cell-division by controlling access to essential nutrients (e.g. nitrogen and phosphorus) (28), which has been previously hypothesized for the Acantharia-*Phaeocystis* symbiosis. We therefore investigated the expression level of marker genes for nutrient deprivation in microalgae: alkaline phosphatase genes for phosphorus limitation (29–31) and nitrate transporter genes for nitrogen limitation (28, 32, 33). Alkaline phosphatase genes (*phoA* and *ehap1*-like) that were expressed by free-living cells cultured in nutrient replete conditions were generally not expressed at detectable levels in symbionts, and the nitrate transporter gene (*Nrt*) was significantly downregulated in symbiotic cells compared to free-living cells (Fig. 2). Hence, symbiotic *Phaeocystis* in the host microhabitat does not appear to be limited by these major essential nutrients, despite very low nutrient availability in the waters from which hosts were collected. This is further supported by the increased nucleolar volume observed in symbiotic cells (Fig. S1), which reflects high rDNA transcription rates (34) and thus increased ribosome production and protein translation—that are processes typically reduced under N-limitation (35). Similarly, the Translation, Translation initiation, and Protein folding GO terms, as well as the Ribosome KEGG pathway, were enriched among the significantly upregulated genes in symbiosis (Fig. S2, Dataset S1). Furthermore, the assimilatory nitrite reductase gene (*NirA*) was downregulated in symbiosis. In contrast, genes responsible for ammonia assimilation (*GS-GOGAT*, *CPS1*) were upregulated (Fig. 2D), indicating that ammonium may be readily available to symbionts, as has been shown in several other photosymbioses (36–38). Together, these results suggest that acantharian hosts may rely on a nutrient-independent mechanism to inhibit symbiont cell division and manage intracellular symbiont populations, a strategy that would give hosts finer control over symbionts while ensuring maximal symbiont productivity.

### Enhanced photosynthesis and carbon fixation in symbiosis

Free-living, flagellated *Phaeocystis* cells usually have two plastids (Figs. 3A and S3). In symbiotic cells, there is a proliferation of enlarged plastids (18), and in this study, we observed 4–65 plastids in individual symbiotic *Phaeocystis* cells. 3D reconstructions allowed us to further determine that plastids occupied 42–62% of the total volume of symbiotic cells, compared to 31% in free-living cells, and that both the total volume and surface area of the photosynthetic machinery expanded up to app. 100 fold (Figs. 1 and 3, Dataset S1). In single-celled algae, plastid division is typically synchronized with cell division and is initiated by expression of nuclear-encoded plastid division genes during S-phase of the cell cycle (39). Several of these genes (*FtsZ*, *DRP5B*, and *PDR1*) were expressed at similar levels in symbiotic and free-living cells (Fig. S4), despite cell-cycle genes being significantly downregulated in symbiosis. Moreover, 3D reconstructions of symbiotic cells revealed several plastids in the process of dividing (Fig. S4). Continued plastid division without consequent cell division leads to the accumulation of plastids and indicates that plastid division has become decoupled from cell division in symbiotic *Phaeocystis*. Symbiotic cells may therefore be arrested in S-phase, thus allowing plastids to proliferate and explaining the build-up of chromatin, but ultimately keeping symbiont population density constrained.

**Figure 3.**
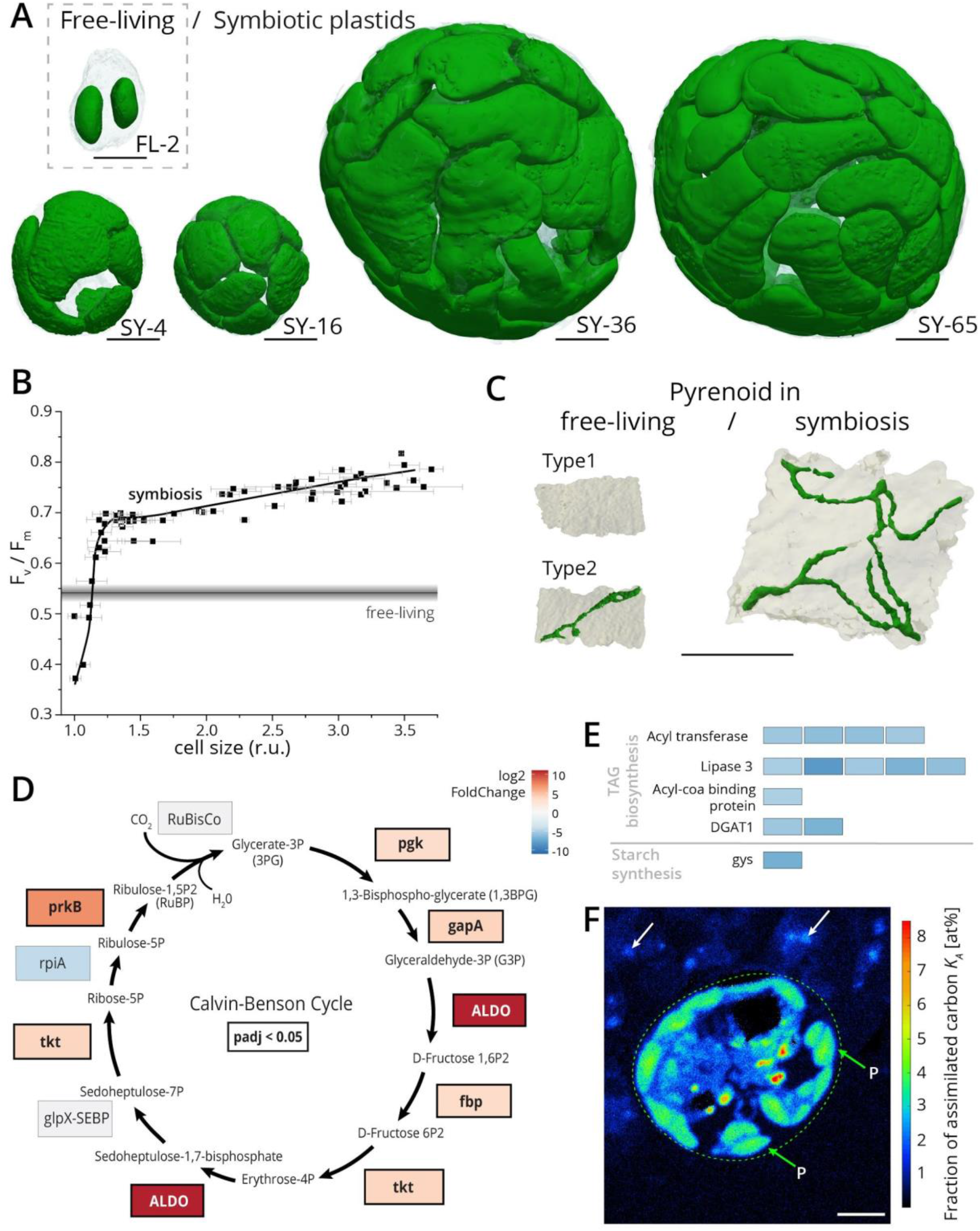
Morphological transformation of the photosynthetic machinery in symbiosis and associated physiological and gene expression activity. **A)** Multiplication of plastids in *Phaeocystis* cells unveiled by FIB-SEM from two in free-living stage (top left) to up to 65 plastids (Sy-65) in symbiosis (see also Fig. S3 and S4). Scale bar: 2 μm. **B)** Single-cell chlorophyll fluorescence imaging showing that the photosynthetic capacity (F_v_/F_m_ parameter) changes as a function of the relative cell size of symbiotic *Phaeocystis* (symbols). The black line represents the mean F_v_/F_m_ value in free-living cells (± s.d., grey lines). **C)** Architecture and organization of pyrenoids in plastids of free-living (two types: with and without tubule) and symbiotic *Phaeocystis*, showing the increased volume of the pyrenoid in symbiosis with multiple thylakoids crossing the pyrenoid (dark green tubules). Scale bar: 1 μm. (See also Fig. S5). **D-E)** Differential expression results for genes encoding enzymes of the Calvin-Benson Cycle (D) and proteins involved in storage molecule biosynthesis (E). The color scale indicates log2 fold-change in symbiosis so that positive values (red) represent upregulation in symbiotic cells and negative values (blue) represent downregulation in symbiotic cells. Significantly differentially expressed genes (false discovery rate adjusted *p*-value, padj < 0.05) are highlighted with bolded gene names and black boxes in D; all results were significant in E. When several isoforms were expressed for a single gene (Acyl transferase, Ligase 3, and DGAT1), the log2 fold-change is shown for each isoform. TAG stands for triacyglycerol. **F)** Single Isotope Probing-NanoSIMS-derived map of carbon relative assimilation (Ka, (70)) showing the fraction of carbon (relative to its initial content) assimilated during 5 hours of incubation with ^13^C-labelled bicarbonate in one symbiotic *Phaeocystis* cell (mainly allocated to plastids) and transferred to the host cytoplasm in specific cellular locations (See also Fig. S6). Green arrows indicate the plastids (P) of symbiotic microalgae (surrounded by the dashed circle) showing about 5 at% of relative carbon assimilation. White arrows indicate host areas revealing about 1–3 at% of the assimilated carbon fraction. Scale bar: 3 μm.

In plants, plastid proliferation increases photosynthesis more efficiently than plastid enlargement and is recognized as a means of increasing surface exchange, particularly for CO_2_ diffusion (40). Here, we found that the photosynthesis GO term and KEGG pathway were both enriched among genes upregulated in symbiotic *Phaeocystis* cells (Fig. S2). To further test whether morphological and transcriptional changes in symbiotic cells enhanced photosynthesis, we assessed the relationship between cell size (a proxy for number of chloroplasts) and photosynthetic efficiency by analyzing photosynthetic parameters *in vivo* at the single-cell level. We found that photosynthetic capacity increased rapidly with cell size until size had doubled, at which point, further increase in photosynthetic capacity was limited (Fig. 3B). This suggests that increased cell size, and therefore expanded photosynthetic machinery, enhances photosynthesis in *Phaeocystis*. The presence of small and large algal cells within a host indicates that *Phaeocystis* cells with different photosynthetic capacity coexist (F_v_/F_m_ ranging from 0.4 to 0.78). The smallest symbiotic algae had F_v_/F_m_ values lower than those measured in free-living cells (0.54 ± 0.02). These algae could represent an early stage during the establishment of symbiosis, in which photosynthesis is transiently repressed.

Expansion of the photosynthetic machinery in symbiotic *Phaeocystis* should be accompanied by increased carbon fixation and production of organic compounds. Carbon uptake was measured in free-living *Phaeocystis* cells and symbiotic *Phaeocystis* within their host incubated with ^13^C-bicarbonate for one hour. We found that symbiotic *Phaeocystis* cells took up approximately 150 times more ^13^C (0.70 ± 0.19 pg of ^13^C.cell^−1^) than free-living cells (0.0043 ± 0.0004 pg of ^13^C.cell^−1^) (Dataset S2). Consistent with this, nearly all nuclear-encoded genes for Calvin-Benson cycle enzymes were upregulated in symbiosis (Fig. 3D). To further explain enhanced carbon fixation in symbiotic *Phaeocystis*, we resolved the internal organization of pyrenoids within plastids. Pyrenoids contain the majority of the Rubisco enzyme in plastids and separate it from the surrounding, more basic stroma to maximize the enzyme’s efficiency and, therefore, the cell’s carbon fixation potential (41). We found that the volume of pyrenoids increased 15 fold from 0.08 ± 0.02 μm^3^ in free-living cells (n = 27 plastids, 14 cells) to 1.2 ± 0.6 μm^3^ in symbionts (n = 74 plastids, 7 cells) and that they occupied a larger proportion of plastid volume (13 ± 2.8% in symbiotic cells compared to 5 ± 1.6% in free-living cells, Fig. 3C, Fig. S5, Dataset S1). A consequence of enlarged pyrenoids in symbiosis is that their surface area:volume ratio inherently decreases (Fig. S5), potentially increasing the diffusion barrier for gas and metabolite exchange. Thylakoid tubules deliver CO_2_ to Rubisco and transport glycerate-3-phosphate product to the stroma, where the remainder of the Calvin-Benson cycles occurs (42, 43). In symbiosis, pyrenoids were always crossed by multiple tubules, whereas pyrenoids in free-living cells either lacked tubules (type 1) or were crossed by one small tubule (type 2) (Fig. 3C). When present, tubules in the pyrenoids of free-living cells were 9.5 times less voluminous than those in symbiotic cells. We hypothesize that multiple tubules in the pyrenoids counterbalance the lower surface area: volume ratio in symbiosis and maintain optimal delivery of CO_2_. These fine-scale changes in pyrenoid structure, alongside upregulation of Calvin-Benson cycle genes, explain the mechanisms underpinning enhanced carbon fixation in symbiotic *Phaeocystis*.

Symbionts may themselves take advantage of the additional organic carbon produced from enhanced carbon fixation—either immediately or after synthesizing storage molecules—or they could transfer the additional fixed carbon to hosts. Based on the transcriptome analyses, symbiotic cells do not seem to be storing more fixed carbon than free-living cells. Key genes for the biosynthesis of triacylglycerol (TAG)—the preferred storage molecule among microalgae in oligotrophic regions (44)—were downregulated in symbiosis, including those for acyl-CoA binding proteins and acyltransferases that are involved in the production of TAG precursors (e.g. phosphatidic acid) and diacylglycerol transferase (DGAT), which performs the terminal step in TAG synthesis (Fig. 3E). Likewise, starch synthase was significantly downregulated in symbiosis (Fig. 3E). Symbiotic cells may store less carbon than free-living cells and instead produce ATP to meet energetic requirements (plastid division, protein synthesis) and relinquish surplus photosynthate to hosts. NanoSIMS analyses incorporating a 5-hour incubation with ^13^C-labelled bicarbonate on four algal cells from two distinct hosts showed that symbiotic *Phaeocystis* cells mainly allocated carbon to their multiple plastids and small vacuoles (Fig. 3F, Fig. S6). These analyses also demonstrated that photosynthetically fixed carbon is transferred to the host cell (Fig. 3F), thus providing direct evidence that the host benefits from the boosted primary production of its symbiotic microalgae.

### Transformation of the mitochondria into a reticulated fine network connected to plastids

Like plastids, mitochondria underwent an extensive expansion in symbiosis, forming a well-developed fine reticular network with mitochondrial volume and surface area increasing up to 52 and 47 fold, respectively (Fig. 4A, Dataset S1). Such reticular and fused networks are characteristic of actively respiring cells in eukaryotes (45). Yet, in relation to cell volume, the contribution of mitochondria was relatively constant (from 5.3 ± 1.5% in free-living to 3.3 ± 0.6% in symbiosis) (Fig. 1C and Dataset S1). At the sub-organelle level, the total volume of mitochondrial cristae (i.e. membrane invaginations, which are respiratory units), increased by up to 38-fold in symbiosis, but volume occupancy was similar in both stages (17.4 ± 3.9% in free-living and 16.3 ± 2.1% in symbiosis) (Fig. S7). The GO terms for Respiration and Oxidative phosphorylation were enriched among genes downregulated in symbiosis (Fig. S2) and the majority of genes in the TCA cycle and respiratory oxidative phosphorylation KEGG pathways were downregulated in symbiosis (Fig. S8). The morphology of the mitochondria and downregulation of respiratory pathways indicates that symbiotic cells may not respire at a higher rate than free-living cells. The mitochondria may, therefore, have a different primary role in symbiotic cells.

**Figure 4.**
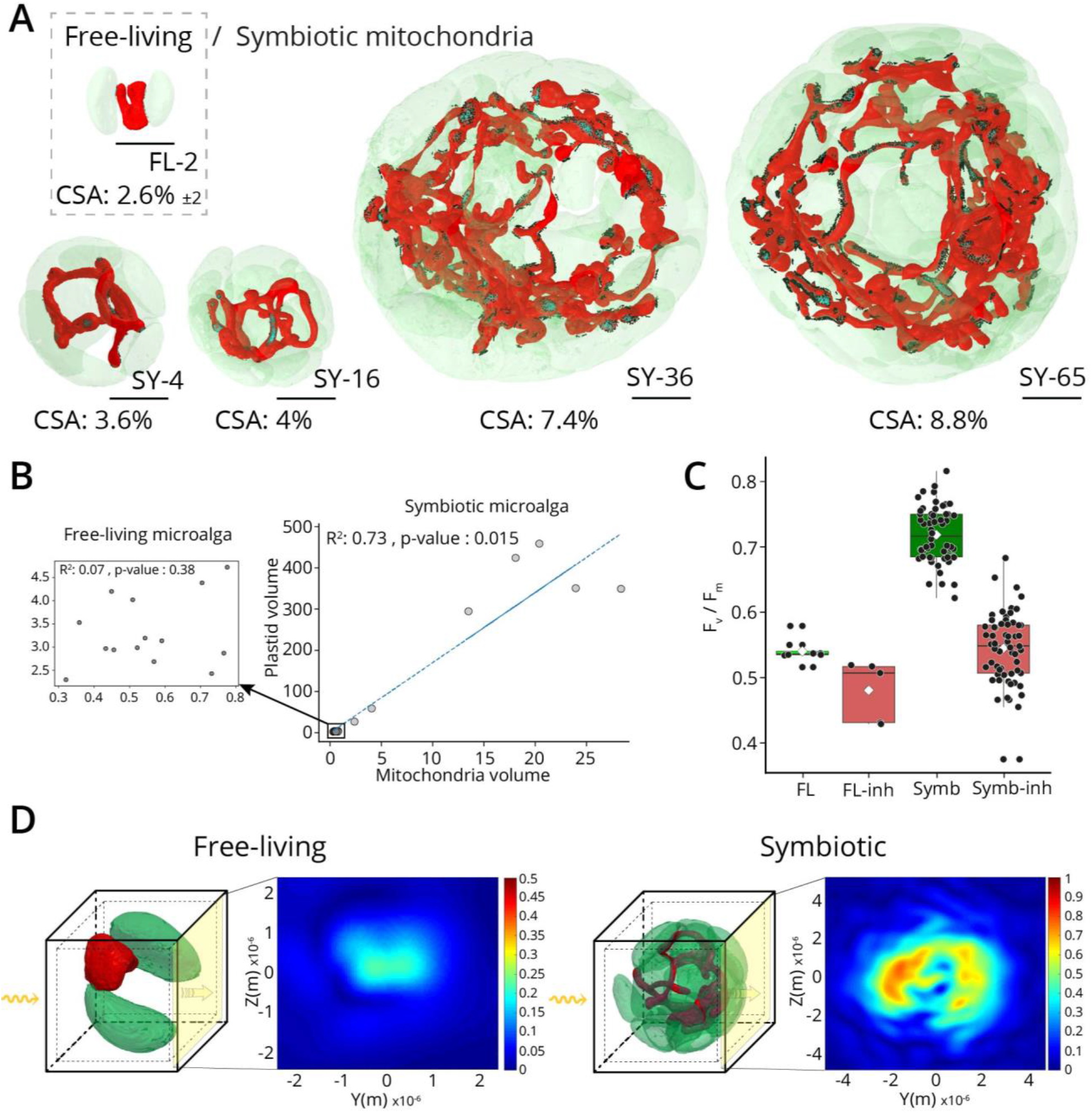
Expansion of the reticulate mitochondria in symbiosis and its interaction with plastids and role in light capture. **A)** Extension of the mitochondria (red) in symbiotic *Phaeocystis* cell towards a fine reticular network in contact with plastids. A free-living *Phaeocystis* cell (top left) and four symbiotic *Phaeocystis* cells (Sy) are represented with their 4, 16, 36 and 65 plastids in transparency. Contact surface area (≤ 50 nm, CSA) between the mitochondria and the plastids have been quantified and represented in dark green colors. The mitochondrial surface area in contact with plastids (CSA) increased from 2.6 ± 2% (n = 14) in free-living to up to 8.79% in symbiosis (Sy-65). (See also Fig. S7). Scale bar: 2 μm. **B)** Scatter plot showing the correlation between the volume of the plastid and the volume of the mitochondria in 20 free-living cells (left insert) and in seven symbiotic *Phaeocystis* cells. **C)** Box plot showing the effect of mitochondrial inhibitors (inh: 5 μM Antimycin A and 1 mM Salicylhydroxamic acid-SHAM) on the F_v_/F_m_ parameter of free-living (left) and symbiotic (right) *Phaeocystis* cells. **D)** Finite-difference-time-domain (FDTD) model calculating the distribution of the light scattering cross-section (σ) based on 3D architectures of microalgal cells. Light scattering by free-living cells (left) is forward directed while light scattering by symbiotic cells (right, 16 plastids) is distributed, yielding an enhanced electric field distribution. Note that due to small light scattering by free-living cells, the scale was adjusted for improved visualisation. The outer and inner boxes represent the scattering monitor and the absorption monitor, respectively. Behind the cell, the yellow plane is measuring the transmitted field direction flow (poynting vector). The yellow arrows correspond to the incident plane wave (See also Fig. S9).

Other metabolic functions could provide a functional rationale for the morphological modification of mitochondria in symbiotic *Phaeocystis*. Mitochondrial interaction with plastids is an essential aspect of the bioenergetics of photosynthetic cells (46–48). In symbiotic *Phaeocystis*, mitochondrial volume tended to increase as plastid volume increased (*R*^2^ = 0.74, *p* = 4.8 E10^−11^), whereas this relationship was not observed for free-living cells (*R*^2^ =0.07, *p* = 0.38) (Fig. 4B). We further found that mitochondrial surface area in contact with plastids (≤ 50 nm distance) increased from 2.63 ± 2.1% in free-living (n = 14 cells) to up to 8.8 % in the largest symbiotic cell that was imaged (65 plastids, Fig. 4A, Dataset S1), with mitochondria interacting with multiple plastids in symbiotic cells. This indicates that the expansion of the reticulated mitochondria is likely related to plastid proliferation, where mitochondria may serve to maintain minimal diffusion distance between the two organelles and ensure optimal metabolic exchange (e.g. lipids, ammonium, ATP; (47, 48)). Consistent with this possibility, a brief incubation with mitochondrial inhibitors (Antimycin A, Salicylhydroxamic acid) at concentrations blocking respiration (47) decreased the F_v_/F_m_ in symbiosis to 0.55 ± 0.05 (Fig. 4C). We interpret this observation in terms of a reduction of the electron acceptors between the two photosystems, due to increased cellular reducing power upon blocking consumption of reducing equivalents by the respiratory chain, as shown in other microalgae (49). This effect of inhibitors was less pronounced in free-living cells, indicating less interaction between the two organelles in this life stage. Overall, these results show that mitochondria contribute to the enhanced physiological performance of symbiotic *Phaeocystis*.

Symbiont photosynthesis within multicellular hosts is strongly affected by the light scattering properties of cells (50, 51). At the cellular level, mitochondria are known to be efficient light scatterers (52). We, therefore, tested whether the well-developed mitochondrial network of symbiotic cells could improve light distribution for photosynthesis. We used Finite-Difference-Time-Domain (FDTD) calculations to model the 3D light propagation in free-living (2 plastids) and symbiotic cells (16 plastids) using the 3D architectures of plastids and mitochondria. We observed that the reticular mitochondrial network homogeneously distributed the incident electric field due to its high scattering cross-section and intricate spatial distribution (Fig. 4D). Contrary to scattering in free-living microalgae, which is small and unidirectional with the incident light, the reticular mitochondrial network of symbiotic microalgae increases the photon pathlength within the algal cell (Fig. 4D and Fig. S9), consequently enhancing the chance of light absorption (51). Thus, mitochondria-induced light scattering may contribute to the optimized photosynthetic performance of *Phaeocystis* in symbiosis. This is consistent with earlier measurements, showing that symbiotic cells have improved photosynthetic efficiency under limiting light (the initial slope **α** of the photosynthesis-irradiance curve, (18)) than their free-living counterparts.

### Physical integration of microalgae into the host cell: symbiosome invagination into symbionts

In photosymbioses, hosts phagocytize microalgal cells from the environment and maintain them individually in a vacuole, or symbiosome, where metabolic exchanges take place. For small symbiotic *Phaeocystis* cells, the symbiosome surrounds the microalgal cell as observed in other photosymbioses (53) (Fig. 5A). Remarkably, in larger symbiotic cells (> 31 plastids), an invagination of the symbiosome vacuole into the microalgal cell was observed in different host cells (Fig. 5B). This invagination can represent a volume of up to 139.4 μm^3^ (one-fifth of symbiont volume) with a tendency to increase with symbiont size (Fig. 5C). We predict that symbiosome invagination maintains/optimizes metabolic exchanges with very large symbionts that would otherwise have decreased surface area for exchange. Of note, small vesicles were visible in the symbiosome space around symbionts, as well as within the invaginated symbiosome in larger symbionts (Fig. 5A and 5B). These vesicles could represent a route for transfering photosynthetic products from symbiont to host or metabolites and signaling molecules from host to symbiont. Indeed, free-living *Phaeocystis* are often considered “secretory cells” because they excrete vesicles rich in organic carbon (polysaccharides) into the environment (54). Penetration of the symbiosome represents a profound morphological manipulation and adds compelling evidence for the concept that symbionts are too far changed to return to their free-living form. Thus, the acantharian host hijacks and parasitizes the microalgal cell, ultimately performing “cytoklepty”. Cytoklepty can be defined as an evolutionarily one-sided endosymbiosis, in which the host captures and physiologically exploits endosymbionts, which eventually die and must be replaced by uptake of new endosymbionts from a wild population.

**Figure 5.**
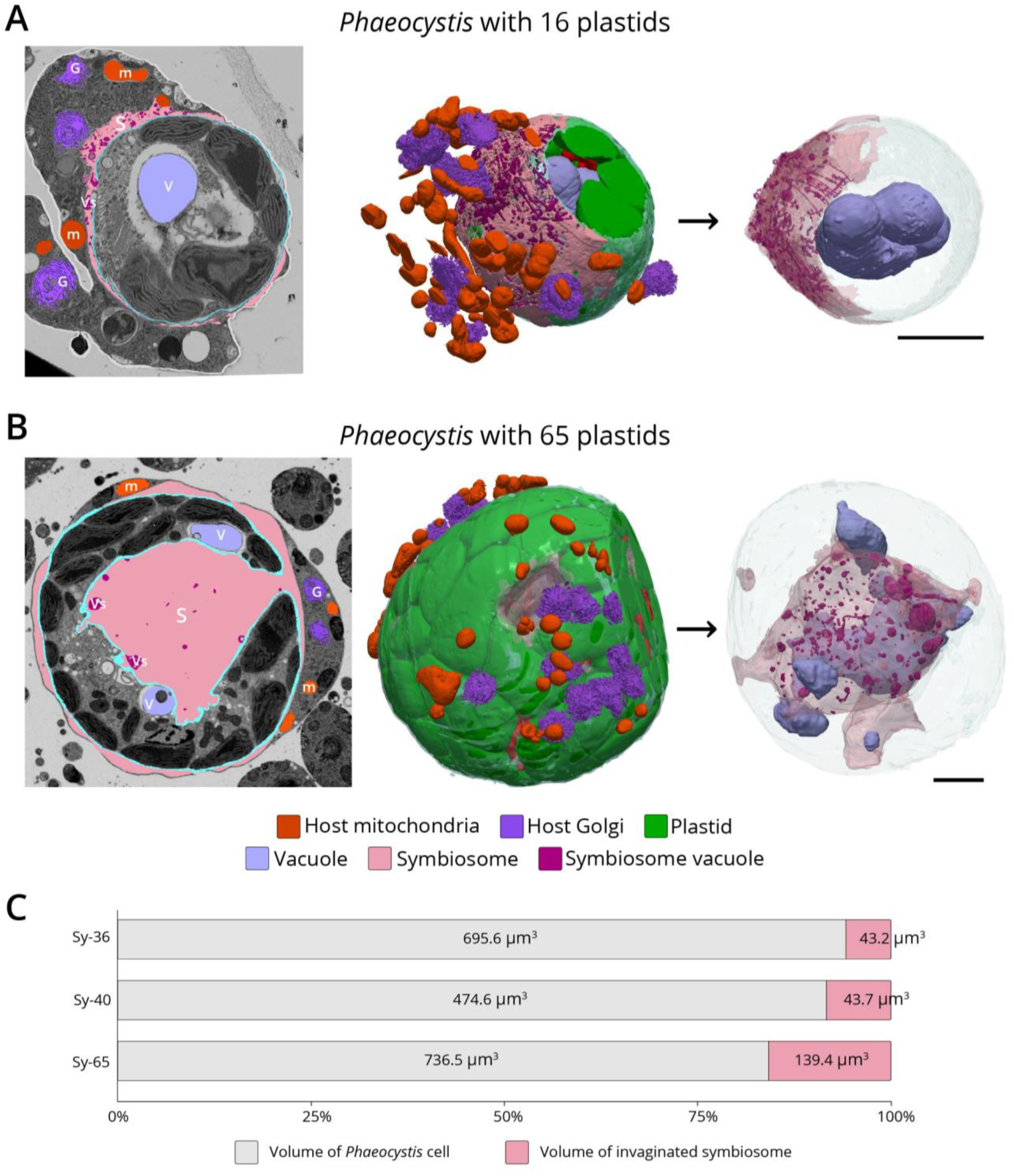
Host-symbiont integration and morphometrics of the symbiosome invagination in large symbiotic microalgae. **A)** 3D reconstruction following FIB-SEM of a symbiotic microalga with 16 plastids surrounded by mitochondria (red) and Golgi apparatus (dark purple) of the host. The different organelles and compartments reconstructed in 3D are highlighted in the FIB-SEM electron micrograph frame (left). 3D reconstruction (right) of a large vacuole in the symbiotic microalga (light purple), which is surrounded by a symbiosome (pink) containing small vesicles (dark red). Scale bar: 2 μm. **B)** 3D reconstruction of a large symbiotic microalga with 65 plastids surrounded by mitochondria (m; red) and Golgi apparatus (G; dark purple) of the host. The different organelles and compartments reconstructed in 3D are highlighted in the FIB-SEM electron micrograph frame (left). In large symbionts (> 31 plastids), there is an invagination of the symbiosome (S; pink) in the algal cell. Note the presence of small vesicles (Vs; dark red) in the symbiosome and large vacuoles (V; light purple) close to the symbiosome. Scale bar: 2 μm. **C)** Cell volume of different symbiotic microalgae with 36, 40 and 65 plastids and the associated invaginated symbiosome that increases in volume.

### Conclusions

The combination of nanoscale imaging with single-cell transcriptomics in this study illuminates the drastic morpho-genetic manipulation of endosymbiotic microalgae involved in a globally distributed and ecologically relevant photosymbiosis. Acantharian hosts prevent *Phaeocystis* cell division, leading to a complete cellular overhaul that improves symbiont bioenergetic performance, enhances photosynthesis and carbon fixation, and ultimately results in substantial organic carbon production and transfer to hosts. More specifically, there is a structural remodeling of the bioenergetic machinery at multiple scales within the algal cell: plastids proliferate and pyrenoids develop with an immersed thylakoid network, which is accompanied by an extension of reticulated mitochondria that maintains contact with plastids and optimizes light distribution within the algal cell. These modifications are clearly beneficial to single-celled planktonic hosts, which gain an additional energy source and avoid symbiont overgrowth. The observation that host symbiosomes intrude into symbiont cells is unique among marine photosymbioses and provides new evidence supporting the idea that the metamorphosis of *Phaeocystis* is unidirectional, making the relationship an evolutionary dead-end for symbionts. Such extreme manipulation of microalgae suggests that this relationship, named cytoklepty, may represent a first step toward a more complete integration of endosymbionts.

Aspects of the symbiont remodeling observed here, such as hypertrophy of the photosynthetic machinery, are also observed in the *Paulinella chromatophora* endosymbiosis, the most recent known example of primary plastid acquisition (55), and in the *Hatena* endosymbiosis, a contemporary example of secondary endosymbiosis in progress (56). Expansion of the photosynthetic machinery may, therefore, represent a common stepping stone towards plastid acquisition. Enlarged nuclei and increased genome ploidy, observed here and in several examples of kleptoplastidy, may represent another commonality in plastid acquisition that serves to support photosynthetic expansion (57, 58).

The uncoupling between cell cycle progression and plastid division in symbiotic *Phaeocystis* suggests that the plastid-to-nucleus signal required to continue cell division is inhibited and deciphering the mechanism involved warrants further attention. Canonical cell-signalling pathways are a promising target for future work, as they have been implicated in mediating other endosymbioses as well as the morphological transformation between flagellate and colonial *Phaeocystis* cells (59, 60). As this cytoklepty shares important similarities with kleptoplastidy and may represent an early intermediary step between photosymbiosis and more permanent incorporation of the photosynthetic machinery, further elucidation of the mechanisms involved should pave the way to a more complete understanding of how eukaryotes acquire new organelles.

## Materials and methods

### Sampling and preparation for electron microscopy

Symbiotic acantharians harboring intracellular microalgal cells (*Phaeocystis cordata*) were collected from surface waters as in (18) (Mediterranean Sea, Villefranche-sur-Mer, France). After collection, individual cells were isolated under a microscope with a micropipette, rapidly transferred into filtered natural seawater, and maintained in the same controlled light (100 μmol photons m^−2^s^−1^) and temperature (20°C) conditions as the free-living stage. In parallel, cultures of the haptophyte *Phaeocystis cordata* (the symbiont of Acantharia in the Mediterranean Sea) (1) (strain RCC1383 from the Roscoff Culture Collection) were maintained at 20°C in K5 culture medium and at 100 μmol photons m^−2^s^−1^.

Sample preparation protocols were adapted from (18) to optimize the contrast for 3D electron microscopy imaging and facilitate pixel classification. Symbiotic acantharians (with algal endosymbionts) and free-living *Phaeocystis cordata* in culture were cryofixed using high-pressure freezing (HPM100, Leica), in which, cells were subjected to a pressure of 210 MPa at 196°C, followed by freeze-substitution (EM ASF2, Leica). Prior to cryo-fixation, the microalgal cultures were concentrated by gentle centrifugation for 10 min. For freeze-substitution (FS), a mixture of dried acetone, and 2% osmium tetroxide and 0.5% uranyl acetate was used as contrasting agents. The FS machine was programmed as follows: 60–80 h at −90°C, heating rate of 2°C h^−1^ to −60°C (15 h), 10–12 h at −60°C, heating rate of 2°C h^−1^ to −30°C (15 h), and 10–12 h at −30°C, quickly heated to 0°C for 1 h to enhance the staining efficiency of osmium tetroxide and uranyl acetate and then back to −30°C. Cells were then washed in anhydrous acetone for 20 min at 30°C and gradually embedded in anhydrous Araldite (resin). A graded resin/acetone (v/v) series was used (30, 50 and 70% resin) with each step lasting 2 h at increased temperature: 30% resin/acetone bath from −30°C to −10°C, 50% resin/acetone bath from −10°C to 10°C, 70% resin/acetone bath from 10°C to 20°C. Samples were then placed in 100% resin for 8–10 h and in 100% resin with the accelerator BDMA for 8 h at room temperature. Resin polymerization finally occurred at 65°C for 48 h.

### Focused Ion Beam Scanning Electron Microscopy (FIB-SEM)

For FIB-SEM, the sample was trimmed with a 90° diamond knife (Diatome) to expose cells at two surfaces (the imaging surface and the surface perpendicular to the focused ion beam) and optimize acquisition (61). For symbiotic *Phaeocystis cordata*, trimming was targeted towards the periphery of hosts where microalgae were more abundant. After the sample was trimmed, it was mounted onto the edge of an SEM stub (Agar Scientific) using silver conductive epoxy (CircuitWorks) with the trimmed surfaces facing up and towards the edge of the stub. The sample was gold sputter-coated (Quorum Q150RS; 180 s at 30 mA) and placed into the FIB-SEM for acquisition (Crossbeam 540, Carl Zeiss Microscopy GmbH). Once the Region of Interest (ROI) was located in the sample, Atlas3D software (Fibics Inc. and Carl Zeiss Microscopy GmbH) was used to perform sample preparation and 3D acquisitions. First, a 1 μm platinum protective coat was deposited with a 1.5 nA FIB current. The rough trench was then milled to expose the imaging cross-section with a 15 nA FIB current, followed by a polish at 7 nA. The 3D acquisition milling was conducted with a 1.5 nA FIB current. For SEM imaging, the beam was operated at 1.5 kV/700 pA in analytic mode using an EsB detector (1.1 kV grid collector voltage) at a dwell time of 8 μs with no line averaging. The voxel size used throughout acquisitions was 5 or 10 nm. Datasets were initially aligned by the Fiji plugin “Linear Stack Alignment with SIFT” (62) then fine-tuned by AMST (63). Raw electron microscopy data are deposited in EMPIAR, accession code EMPIAR-XXX.

### FIB-SEM Image analysis: segmentation and morphometric analyses

From the aligned FIB-SEM stack, images were binned in Fiji (https://imagej.net/Fiji), and regions of interest representing *Phaeocystis* cells were cropped. Segmentation and 3D reconstruction were performed using the work-flow developed in (64), and geometry measurements were provided using algorithms provided at (https://gitlab.com/clariaddy/stl_statistics and https://gitlab.com/clariaddy/mindist). Briefly, segmentation of organelles and vacuoles of *Phaeocystis* (considered as regions of interest) was carried out with 3DSlicer software (65) (www.slicer.org), using a supervised semi-automatic pixel classification mode (3 to 10 slices automatically segmented for each ROI). Each region was “colored” using paint tools and adjusting the threshold range of pixels values of the images. The Model maker module from 3D slicer was then used to generate corresponding 3D models which were exported in STL format. 3D reconstructed models were imported into the MeshLab software (66) to clean the model and reduce file size by model decimation. Metrics for volumes, surface area, and the area below the minimum distance between meshes were computed using Numpy-STL (https://pypi.org/project/numpy-stl/) and TRIMESH (https://trimsh.org/trimesh.html) python packages. Surfaces and volumes were computed using the discrete mesh geometry, the surface being computed directly from mesh triangles, and volume being obtained from the signed volume of individual tetrahedrons, assuming a watertight model. Proximity distance between organelles was calculated based on the closest points between two triangular meshes. The surface area below the proximity distance was quantified based on (i) the minimal distance between each vertex of the plastid mesh to the mitochondria mesh, and (*ii)* matching surface using face data according to a given distance threshold. A distance threshold ≤ 50 nm was chosen as representative of an interaction between nearby organelles, based on previous morphometric analyses in animal and plant cells (48, 67). Morphometric data are available in Dataset S1.

### Single-cell chlorophyll fluorescence imaging

Single-cell chlorophyll fluorescence was detected with an imaging system (JBeamBio, France), mounted on an optical microscope (CKX 53 Olympus, Japan). The imaging setup consists of a high sensitivity camera (Orca Flash 4.0 LT, Hamamatsu, Japan) equipped with a near-infrared long-pass filter (RG 695 Schott, Germany). Regions of Interest (ROIs) containing host cells were identified using the microscope in transmission mode. The photosynthetic capacity of symbiotic *Phaeocystis* was quantified as the photochemical quantum yield of Photosystem II (F_v_/F_m_), calculated as (F_m_−F_o_)/F_m_ (68). Cells were excited with blue light pulses (l = 470 nm ± 12 nm, duration 260 μs) to evaluate minimum fluorescence emission F_o_. Short green saturating pulses (l = 520 nm ± 20 nm, intensity 3000 μmol photons m^−2^ s^−1^, duration 250 ms) were used to reach maximum fluorescence emission (F_m_). F_m_ was evaluated with the same blue pulses used for F_o_, fired 10 μs after the saturating light was switched off. We used a 20X (NA = 0.45) objective to scan the slits. Single algal cells within a given ROI were imaged separately, with a pixel resolution of 1.7 μm^2^.

### Comparative transcriptomics for symbiotic and free-living Phaeocystis

Individual acantharians (n = 12) were collected for single-host RNA-seq from the western subtropical North Pacific. RNA-seq was also performed with biological culture replicates (n = 3) of *Phaeocystis cordata* CCMP3104 (synonymous to RCC1383) to assemble a *de novo* reference transcriptome and to provide a comparison point for changes in symbiotic gene expression. Raw sequencing reads are available from the NCBI Sequencing Read Archive with accession PRJNA603434. Detailed descriptions of sampling, culturing, RNA extraction, sequencing, transcriptome assembly and annotation, differential expression and functional enrichment testing are presented in the SI Appendix and in Tables S1 and S2 and Fig. S10. All code and data analysis pipelines are available from GitHub (https://maggimars.github.io/PcordataSymbiosisDGE/analysis.html). Genes were considered significantly differentially expressed when the False Discovery Rate adjusted *p*-value (padj) was less than 0.05 and the log2 fold-change was greater than |1|. Likewise, Gene Ontology (GO) and KEGG (Kyoto Encyclopedia of Genes and Genomes) pathways were considered significantly enriched among up- or downregulated genes when the padj from the respective enrichment test was less than 0.05.

### ^13^C bulk enrichment (EA-IRMS) and isotope analysis

Symbiotic acantharia were sampled as described above. Between 54 and 78 individual acantharian cells were pooled together. Two control samples were kept in unspiked filtered seawater (FSW, 0.22 μm) to obtain natural (background) ^13^C cell content and two experimental samples were incubated for one hour in ^13^C-bicarbonate spiked FSW at 20°C under constant light (100 μmol photons m^−2^s^−1^). To start the incubation, H^13^CO_3_ (99%^13^C; Cambridge Isotopes Laboratory Inc.) was added to the FSW (0.2 mM, 10% final concentration). After one hour, acantharian cells were immediately harvested by centrifugation and rinsed three times with FSW and one time with Milli-Q water. All individuals in each sample were then transferred to tin capsules and dried at room temperature for three days.

In parallel, cultures of free-living *Phaeocystis cordata* RCC1383 maintained at 20°C in K5 culture medium and at 100 μmol photons m^−2^s^−1^ were also incubated for 1 hour with ^13^C-bicarbonate spiked K5 medium (0.2 mM, 10% final concentration). As for symbiotic cells, two samples were used as controls to obtain natural (background) ^13^C cell content and were kept in unspiked K5 medium. To harvest cells, cultures were centrifuged and rinsed four times with FSW (0.22 μm). Before free-living were transferred to tin capsules, an aliquot was taken from each sample to count the number of cells per sample (Dataset S2). Tin capsules were dried for one day at 37°C.

Samples were analyzed for ^13^C-enrichment using an Elemental Analyser (Flash 2000, Thermo scientific, Milan, Italy) coupled to an Isotopic Ratio Mass Spectrometer (IRMS Delta V Plus with a Conflo IV Interface, Thermo scientific, Bremen, Germany). Atom% of samples was derived from isotope ratio data and calculated using the Vienna Pee Dee Belemnite standard (*R*VPDB = 0.0112372)

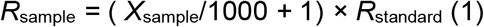

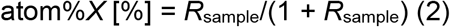

The uptake of ^13^C (*I_isotope_*) in acantharia and *Phaeocystis* cells was obtained from the excess *E* of the ^13^C stable isotope in the sample above background (*E* = atom%*X*_sample_ − atom%*X*_control_), and total carbon content in the sample or cell.

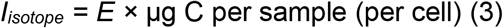

To measure ^13^C uptake of symbiotic *Phaeocystis* cells in Acantharia, we estimated that one acantharian cell harbors an average of 30 cells (29 ± 27, (1)). Note that we also added the values calculated for 60 symbiotic *Phaeocystis* per Acantharia to have conservative estimations of ^13^C-uptake in Acantharia cells (Dataset S2). The uptake of ^13^C (*I_isotope_*, pg. cell^−1^) for symbiotic *Phaeocystis* cells was calculated using these estimations. A short incubation (1 hour) was chosen to reduce the effect of cell respiration and C-compound exudation on measured ^13^C uptake values.

### NanoSIMS imaging and associated sample preparation

Upon collection, symbiotic acantharians were maintained in filtered seawater for 24 hours. Cells were then incubated with ^13^C-labelled bicarbonate (0.4 mM, corresponding to a 16% ^13^C labeling) for five hours and cryofixed as for electron microscopy according to a freeze substitution protocol from (18). Semi-thin sections (200 nm) were placed on boron-doped silicon wafers, coated with a 20 nm Au/Pd (80/20) layer and analyzed with a nanoSIMS 50L (Cameca) at the ProVIS Centre for Chemical Microscopy (UFZ Leipzig). The analysis areas involving symbiotic algal cells were defined from SEM observations on consecutive sections. Before analysis, pre-implantation with 200 pA of 16 keV Cs^+^ ion beam was performed for 15–20 min in 100×100 μm² area to equilibrate the yield of negative secondary ions. Upon analysis, a 16 keV Cs^+^ primary ion beam of 1–2 pA focused to approximately 70 nm was rastered over the sample area between 15×15 μm^2^ and 70×70 μm^2^, with a dwelling time of 2 ms pixel^−1^ in a 512×512 or 1024×1024 pixel pattern, keeping the physical pixel size well below the beam-spot size in order to avoid an ion-beam induced surface roughening. Secondary ions extracted from each pixel of the sample surface were analyzed for their mass to charge (m/z) ratio and counted separately with seven electron multiplier detectors. To resolve isobaric interferences, the mass resolving power (MRP) of the spectrometer was set > 8000 (M/DM) with the entrance slit of 20×140 μm (width × height; nominal size), 200×200 μm aperture, 40×1800 μm exit slits and the energy slit cutting-off 20% of ^12^C^14^N^−^ ions at their high-energy distribution site. Secondary ion species (^16^O^−^, ^12^C_2_^−^, ^12^C^13^C^−^, ^12^C^14^N^−^, ^13^C^14^N^−^, ^31^P^−^, ^32^S^−^) were simultaneously detected in single-ion counting mode. The acquired 40 serial maps of secondary ion count were corrected for lateral drift and accumulated with the Look@NanoSims software (69). Isotope ratios were calculated for each ROI defined with LANS either in automatic thresholding mode or by manual drawing over cell compartments recognized in ion-count/ratio maps (normalized by scans and pixel number) and correlative SEM images. The fraction of assimilated carbon relative to its initial content (relative carbon assimilation, Ka) was derived from the changes in carbon-isotope composition as described in (70).

### Photon propagation model: Finite-difference time-domain simulations

To study the role of light scattering from the 3D mitochondrial network we performed 3D finite-difference time-domain (FDTD) calculations using a Maxwell equation solver (Lumerical FDTD Solutions 8.16). FDTD allows for discretizing the real-space time-domain Maxwell equations onto a regular lattice in time and space with equidistant time steps and cubic voxels on the Yee grid. The propagation of the electromagnetic field is modelled by time stepwise forward integration. For plastids, we assumed that the real part of the refractive index was minimally wavelength dependent (between 1.352–1.364 for 400–700 nm), while the imaginary part of the refractive index (*k*) was governed by the characteristic absorption profile of chlorophyll *a* (71). The real refractive index (*n*) of mitochondria was assumed to be 1.4, which is a moderate estimate for such strongly light scattering structures (52, 72) with a reduced scattering coefficient about two orders of magnitude greater than the absorption coefficient (μs’ > 100*μa) (52). The background *n* of the cell was similar to water (n = 1.34). For each cell type (i.e. free-living and symbiotic) we performed calculations in the presence and absence of mitochondria. Additionally, we replaced the mitochondria with a solid sphere of the same volume, to visualise the beneficial light spreading by the intricate mitochondrial network over a solid sphere with the same refractive index properties. For each simulation, excitation was provided by a plane wave that was incident in either the x, y or z plane. Only one plane is shown as the results were largely independent of the incident plane. The numerical stability of the simulation was ensured by selecting boundary conditions and simulation times (> 150 fs) that confirmed that the electric field in the structure decayed prior to the end of the simulation, such that all of the incident excitation was lost from the grid.

## Acknowledgments

This project received funding from the ATIP-Avenir program, a Défi X-Life grant from CNRS, the LabEx GRAL (ANR-10-LABX-49-01), financed within the University Grenoble Alpes graduate school (Ecoles Universitaires de Recherche) CBH-EUR-GS (ANR-17-EURE-0003). J.D. was supported by the ATIP-Avenir program. C.U. was supported by a joint UGA-ETH Zurich PhD grant in the framework of the “Investissements d’avenir” programme (ANR-15-IDEX-02). M.M.B. was supported by a DC1 graduate fellowship awarded by the Japan Society for the Promotion of Science. D.W. was funded by the Gordon and Betty Moore Foundation and L.S. was supported by the Isaac Newton Trust. G.F. and D.F. received funding from the European Research Council: ERC Chloro-mito (grant no. 833184). This project also received funds from the European Union’s Horizon 2020 research and innovation programme CORBEL under the grant agreement No 654248. We thank the institutes that supported the collection of samples: EMBRC-France and the Laboratoire d’Océanographie de Villefranche-sur-Mer (John Dolan and the marine crew). This work used the platforms of the Grenoble Instruct-ERIC centre (ISBG ; UMS 3518 CNRS-CEA-UGA-EMBL) within the Grenoble Partnership for Structural Biology (PSB), supported by FRISBI (ANR-10-INBS-05-02) and GRAL, financed within the University Grenoble Alpes graduate school (Ecoles Universitaires de Recherche) CBH-EUR-GS (ANR-17-EURE-0003). We thank Guy Schoehn and the electron microscope facility, which is supported by the Auvergne-Rhône-Alpes Region, the Fondation Recherche Medicale (FRM), the fonds FEDER and the GIS-Infrastructures en Biologie Sante et Agronomie (IBISA). We are thankful for the use of the analytical facilities of the Centre for Chemical Microscopy (ProVIS) at UFZ Leipzig, which is supported by European Regional Development Funds (EFRE—Europe funds Saxony) and the Helmholtz Association. Sampling and molecular work for transcriptome analysis was funded by the Marine Biophysics Unit of the Okinawa Institute of Science and Technology (OIST) Graduate University. We thank the captain and crew of the JAMSTEC R/V *Mirai* for their assistance and support in sample collection. Hiromi Watanabe, Dhugal Lindsay, and Yuko Hasagawa were instrumental in organizing and facilitating cruise sampling. Lisa Mesrop and Dave Caron collaborated to optimize single-host RNA-seq. Hiroki Goto and the OIST DNA Sequencing Section performed sequencing and provided guidance. We also thank Gaël Guillou, Benoit Lebreton and the Plateforme de Spectrométrie Isotopique of the La Rochelle University, which is funded by FEDER Poitou-Charentes and the Région Nouvelle Aquitaine (2017-2021).

