## Supplemental Materials for "Cytoklepty in the plankton: a host strategy to optimize the bioenergetic machinery of endosymbiotic algae"

**Transcriptomic analyses of free-living and symbotic stages of the microalga *Phaeocystis***

**Acantharian collection for transcriptomics**

Individual acantharians (n = 12) were collected from 5 sampling stations in the Okinawa Trough (East China Sea) in May and June 2017 during the Japan Agency for Marine-Earth Science and Technology (JAMSTEC) MR17-03C cruise. Plankton samples were collected by passing unfiltered seawater pumped from the sea surface through a 100-µm-mesh-size, hand-held plankton net (Rigo) and were observed under a dissecting microscope. Individual acantharians were transferred by glass micropipette to clean Petri-dishes, rinsed with 0.2-µm-filtered seawater several times, until all visible contaminants were removed, and incubated for 0.5–2 hr for additional self-cleaning. Each acantharian was imaged with inverted light microscopy (Zeiss Primovert) before being transferred to a maximum recovery PCR tube (Axygen) with the smallest possible volume of accompanying seawater. Transfer success was confirmed by microscopy before adding 30 µL of RLT-plus cell-lysis buffer (Qiagen) and immediately flash-freezing in liquid nitrogen. Samples were stored at -80°C until RNA extraction.

***Phaeocystis* culture conditions for transcriptomics**

Three biological replicates of *Phaeocystis cordata* CCMP3104 were prepared for RNA-seq following the methods described in Mars Brisbin et al. (1). Briefly, culture replicates were initiated by inoculating 45 mL of sterile L1 media with 1 mL stock culture in autoclaved glass 50-mL Erlenmeyer flasks. Replicates were placed on a gently rotating twist mixer in a plant growth chamber with cool white fluorescent lamps (CLE-305, TOMY) set to 22°C with light level 4 and a 12 h:12 h day:night ratio. Four days after initiating cultures, 1 mL of each replicate was transferred to 45 mL of sterile L1 media in a clean 50-mL flask and the culturing set-up was repeated. Algal cells were harvested for RNA extraction on the fourth day of the second culture round by filtering entire culture volumes through polytetrafluoroethylene (PTFE) filters (0.45-µm pore-size, Millipore) under gentle vacuum. Filters were immediately flash-frozen in liquid nitrogen and stored at -80°C until RNA extraction.

**RNA extraction and sequencing library preparation**

RNA extractions from single acantharian holobionts were accomplished by modifying methods of Trombetta et al. (2). Samples were thawed over ice, vortexed twice (10 s, speed 7, Vortex-Genie 2), and then incubated at room temperature for 5 min to fully lyse cells. Agencourt RNAClean XP magnetic beads (Beckman Coulter) were added to each sample at a 2.2:1 V:V ratio and fully mixed by pipette prior to a 30-min incubation to bind all RNA to the magnetic beads. After two 80% ethanol washes, RNA was eluted from the beads in 11 µL of a custom elution buffer (10.72 µL nuclease-free water, 0.28 µL RNAase inhibitor) and 10.5 µL of eluted RNA were directly processed following the single-cell protocol for the SMART-seq v4 Ultra Low Input Kit (Clonetech/Takara) with 18 cycles in the primary PCR.

RNA was extracted from *Phaeocystis* cells collected on PTFE filters by following the manufacturer's protocols for the MoBio PowerWater RNA extraction kit (Qiagen) including the optional initial heating step. Extracts were diluted so that 10 ng of RNA were used from each sample as input with the SMART-seq v4 Ultra Low Input RNA Kit. Each *Phaeocystis* sample was additionally supplemented with 2 µL of a 1:10,000 dilution of External RNA Controls Consortium (ERCC) spike-in mix one (Ambion) as an internal quality control. ERCC spike-in was only added to *Phaeocystis* libraries, and not single holobiont libraries, because RNA concentrations in single-holobiont libraries were extremely low and the spike-in would, therefore, utilize too much of the sequencing resources for those samples. The SMART-seq total RNA protocol was then followed with 12 cycles in the primary PCR.

The quality and concentration of cDNA from acantharian holobionts and *Phaeocystis* cultures were assessed with Qubit and Bioanalyzer high sensitivity DNA assays before continuing with the manufacturer's protocols for the Nextera XT DNA Library Prep Kit (Illumina). Twelve acantharian holobiont and three *Phaeocystis* culture cDNA libraries were submitted to the Okinawa Institute of Science and Technology DNA Sequencing Section. Sequencing libraries were pooled and sequenced with 2x150bp paired-end sequencing chemistry on an Illumina HiSeq4000.

Sequencing for this project produced over 2.9 billion read pairs with 78–350 million read pairs per sample. The sequencing data are available from the NCBI Sequence Read Archive (SRA) with accession number PRJNA603434. Following quality filtering with Trimmomatic, 2.5 billion read pairs remained, with 62–305 million read pairs per sample. Reads mapping to ERCC reference sequences (3) were counted with RSEM software for each culture sample (4) and counts were further analyzed in the R statistical environment (5). To assess the relationship between ERCC sequence read counts and their original concentrations in the ERCC internal quality standard, the log of the observed FPKM (Fragments Per Kilobase per Million reads) was plotted against the log of the initial concentrations for each standard sequence. Linear regressions were fit for each sample and $R^2$ values ranged from 0.883–0.926 (Fig. S10). The strong relationship between observed FPKM and initial concentration for ERCC sequences indicates minimal bias was introduced during PCR amplification, library preparation, and sequencing.

### ***Phaeocystis*** reference transcriptome assembly and annotation

Adapters were trimmed from sequencing reads and low-quality reads were filtered and discarded with trimmomatic software (6). Reads mapping to ERCC reference sequences were removed to prevent internal standard sequences from being included in the transcriptome assembly. The pre-processed *Phaeocystis cordata* culture libraries were then assembled into a *de novo* transcriptome using the Trinity software v2.8.4 (7), which was then deduplicated by removing contigs with 95% similarity using CD-HIT-EST (8). While the SMART-seq kit employs poly-A priming to target eukaryotic mRNA and reduce the amount of ribosomal and bacterial RNA present in sequencing libraries, some RNA deriving from these sources remain, especially in low-input samples. Sequences of bacterial origin were bioinformatically removed from the transcriptome by performing a blastn query against the NCBI nr-nt nucleotide database (downloaded March 2018, ncbi-blast v2.6.0+, (9)) and parsing results to identify and remove bacterial contigs.

The *Phaeocystis cordata* transcriptome was annotated using two different databases and functional annotation methods: Pfam (10) and the Kyoto Encyclopedia of Genes and Genomes (KEGG) (11). Pfam annotation was accomplished with the dammit software, which wraps Transdecoder to translate transcriptome contigs to the longest possible amino acid sequence (12) and HMMER to assign Pfam protein homologs to sequences (13). After discarding annotations with e-values greater than 1E-5, the Pfam annotation with the lowest e-value was selected for each contig. The Pfam annotations were matched to corresponding Gene Ontology (GO) terms using the Gene Ontology Consortium's Pfam2GO mapping file (geneontology.org/external2go/pfam2go, version 07/14/2018, (14). KEGG Orthology (KO) annotation was performed using the GhostKOALA tool with the Transdecoder translated amino acid sequences (kegg.jp/ghostkoala, 05/21/2019, (11). The KEGG Orthology numbers were then used to access the KEGG API (kegg.jp/kegg/rest/keggapi.html, July 2019) and determine the KEGG pathways to which transcriptome contigs belonged. Additional annotations for genes of interest were acquired by blasting the *Phaeocystis* transcriptomes against the *Emiliania huxleyi* reference genome (15). Assembly and annotation results are summarized in Table S2. Final assemblies were assessed for completeness by determining how many eukaryotic and protistan Benchmarking Universal Single-

Copy Orthologs (BUSCO v3, (16)) were present in the transcriptomes (Fig. S10). The *P. cordata* CCMP3104 BUSCO scores were compared with scores for the *P. cordata* RCC1383 transcriptome that was assembled as part of the Marine Microbial Eukaryote Transcriptome Sequencing Project (MMETSP) (17).

### Differential gene expression analysis

Quality filtered sequences from each of the twelve acantharians and the three *P. cordata* culture replicates were mapped to the reference transcriptome assembled for this study. All read mapping and quantifying steps were accomplished with the Salmon software (18). Counts for each sample were imported into the R statistical environment with tximport (19). Differential gene expression testing between symbiotic and free-living replicates was performed with the DESeq function in the Bioconductor package DESeq2 (20). Genes that were differentially expressed with a False Discovery Rate (FDR) adjusted *p*-value (padj) less than 0.05 and a log2 fold change > |1| were considered statistically significant and included in gene set enrichment testing. The majority of genes were constitutively expressed (35,829/41,629). Among genes that were significantly differentially expressed in symbiosis, 5,211 genes were downregulated and 589 were upregulated (Fig. S10). A principal component analysis (PCA) on the log normalized counts showed that the majority of the variance among samples (63%) could be explained by symbiosis (Fig. S10)

### Functional enrichment testing

GO term enrichment among significantly up- and downregulated genes was determined with a hypergeometric test in the R package GOstats (21). GOstats accommodates user-defined GO annotations, which are necessary when studying non-model organisms like *Phaeocystis*. Enrichment of KEGG pathways among up- and downregulated genes was tested for by applying linear model analysis with the kegga function from the Bioconductor package edgeR (22). GO terms and KEGG pathways were considered significantly enriched when the statistical test resulted in a *p*-value of less than 0.05. GO terms were further categorized and visualized with REVIGO software (23) and GO terms enriched in up- and downregulated genes were plotted as treemaps (Fig. S2). Consistent with the lower annotation rates for KEGG terms, less KEGG pathways were enriched among up- and downregulated genes in symbiotic *Phaeocystis* than were GO terms. Ten KEGG pathways were enriched among genes upregulated in symbiosis and sixteen pathways were enriched among downregulated genes (Table S1). KEGG pathways were further visualized with the pathview R package (24).

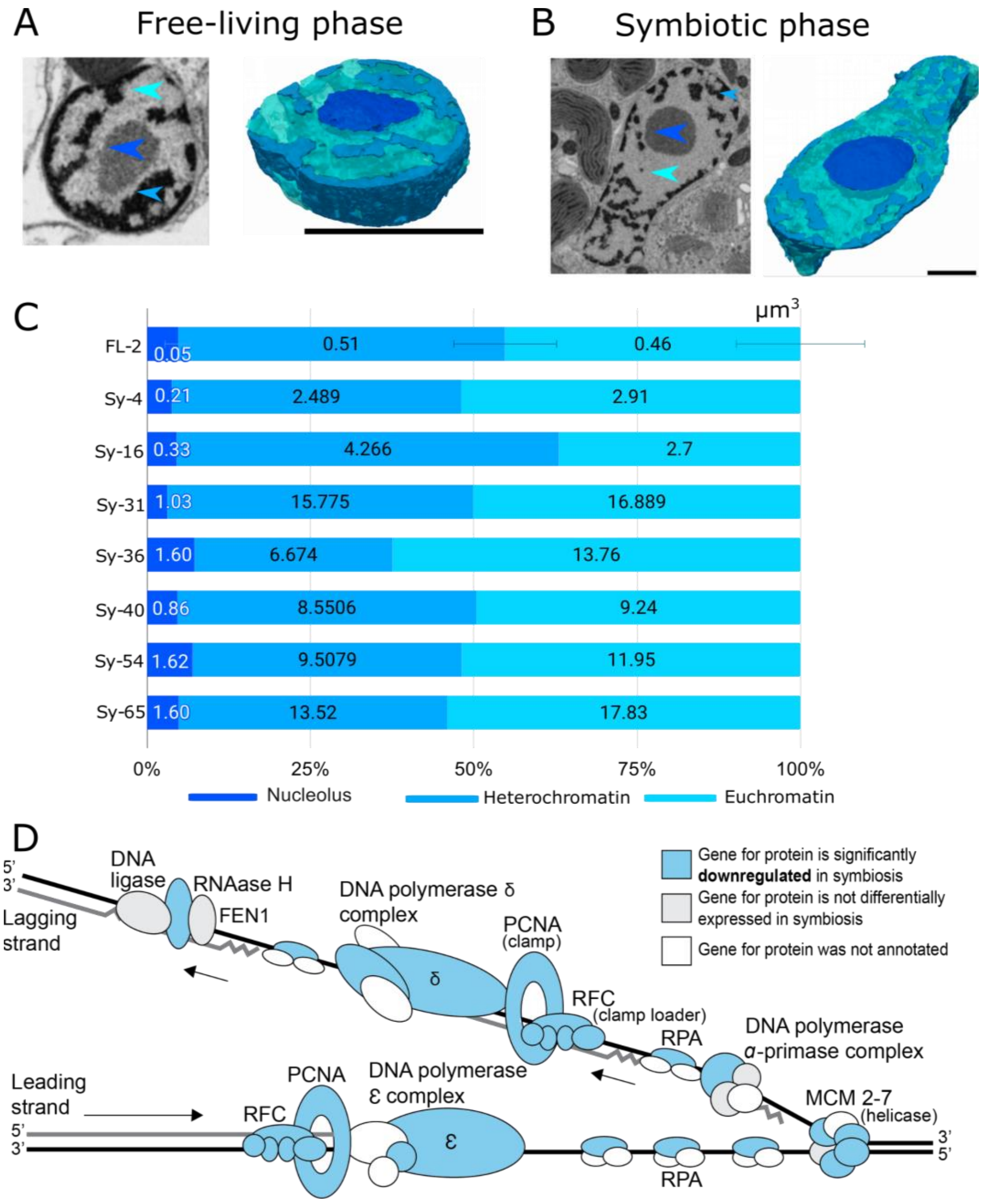

**Fig. S1. Nucleus morphometrics of the microalga *Phaeocystis* and expression of genes involved in DNA replication.** Nucleus morphology in free-living **(A)** and symbiotic stage **(B)** of the microalga *Phaeocystis* unveiled by FIB-SEM imaging. Inside the nucleus, different nuclear compartments were segmented, such as the nucleolus (dark blue), euchromatin (light blue) and heterochromatin (blue). Scale bar: 1 µm. **(C)** Occupancy (% of the cell volume) of the nucleolus, heterochromatin and euchromatin in free-living cells (n = 14), and different symbiotic *Phaeocystis* cells with 4, 16, 31, 36, 40, 54, 65 plastids. The volume ($\mu m^3$) of each of these 3 nuclear compartments are indicated in the corresponding bar charts. **(D)** DNA replication genes are significantly downregulated in symbiotic *Phaeocystis*. Differential expression of genes in the DNA replication KEGG reference pathway (ko03030) Eukaryotic DNA replication complex. Proteins for which genes were significantly downregulated in symbiosis ($\log2FC < -1$ and $padj \leq 0.05$) are solid blue. Genes for subunits in grey were not significantly differentially expressed in symbiosis. If genes for the protein were not annotated in the transcriptome, the subunit is white. The majority of genes in this pathway were downregulated and no genes for DNA replication complex subunits were significantly upregulated in symbiosis.

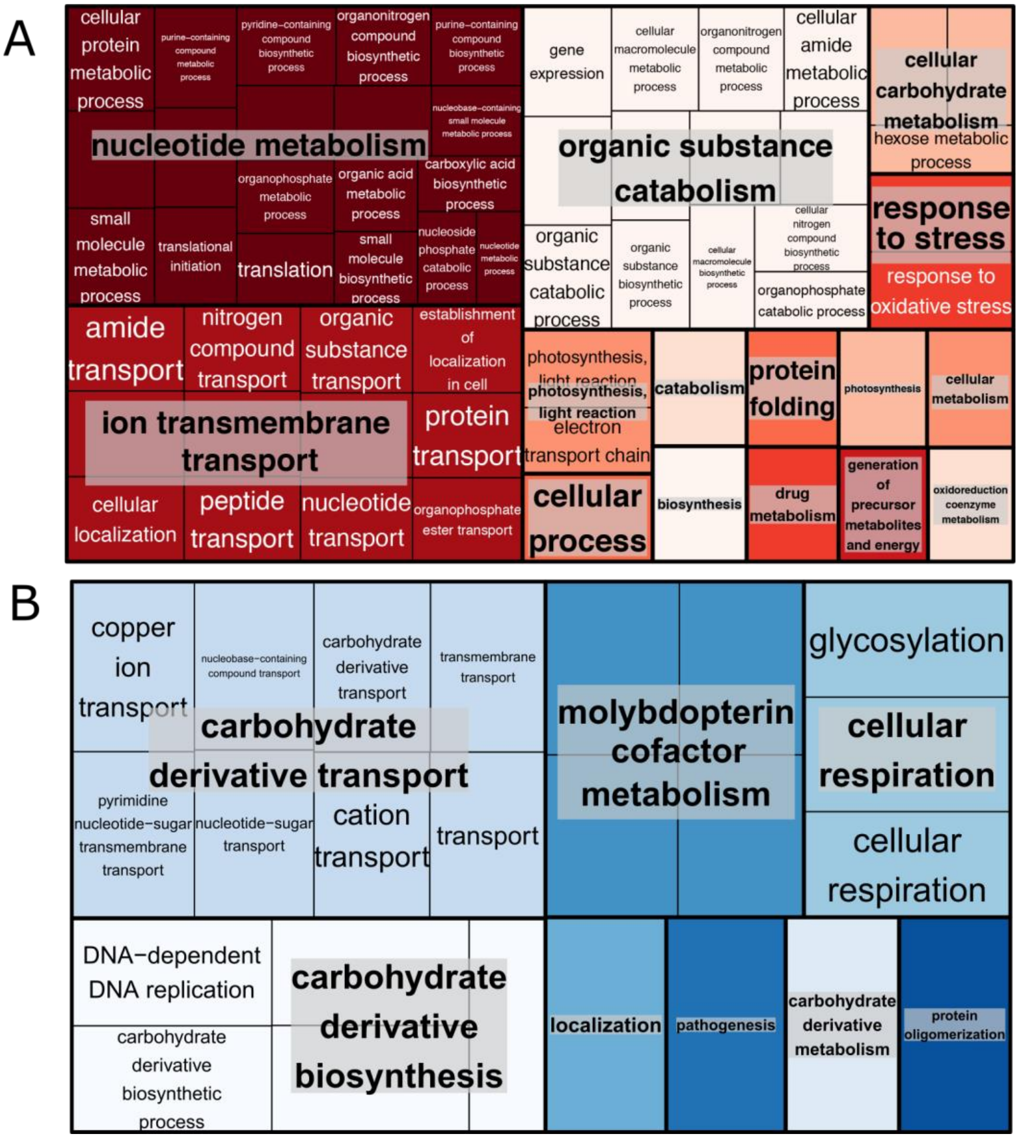

**Fig. S2. Functional enrichment results for *Phaeocystis* genes that were differentially expressed in symbiosis. (A)** Treemap illustrating Gene Ontology (GO) terms significantly enriched among genes that were upregulated in symbiosis (p-values < 0.05, hypergeometric test with GOstats package). There were 139 biological process (BP) GO terms enriched among the 589 *Phaeocystis* genes that were upregulated in symbiosis. Full results from enrichment testing with upregulated genes are available from https://tinyurl.com/cordata-UPregulated-GO-term. The full list of enriched GO terms was processed to reduce redundancy and categorize terms before visualizing results as a treemap. **(B)** Treemap illustrating GO terms enriched among genes that were significantly downregulated in symbiosis. Thirty-one BP GO terms were enriched among the 5,211 *Phaeocystis* genes downregulated in symbiosis. Full results from enrichment testing with downregulated genes are available from https://tinyurl.com/cordata-DOWNregulated-GO-term.

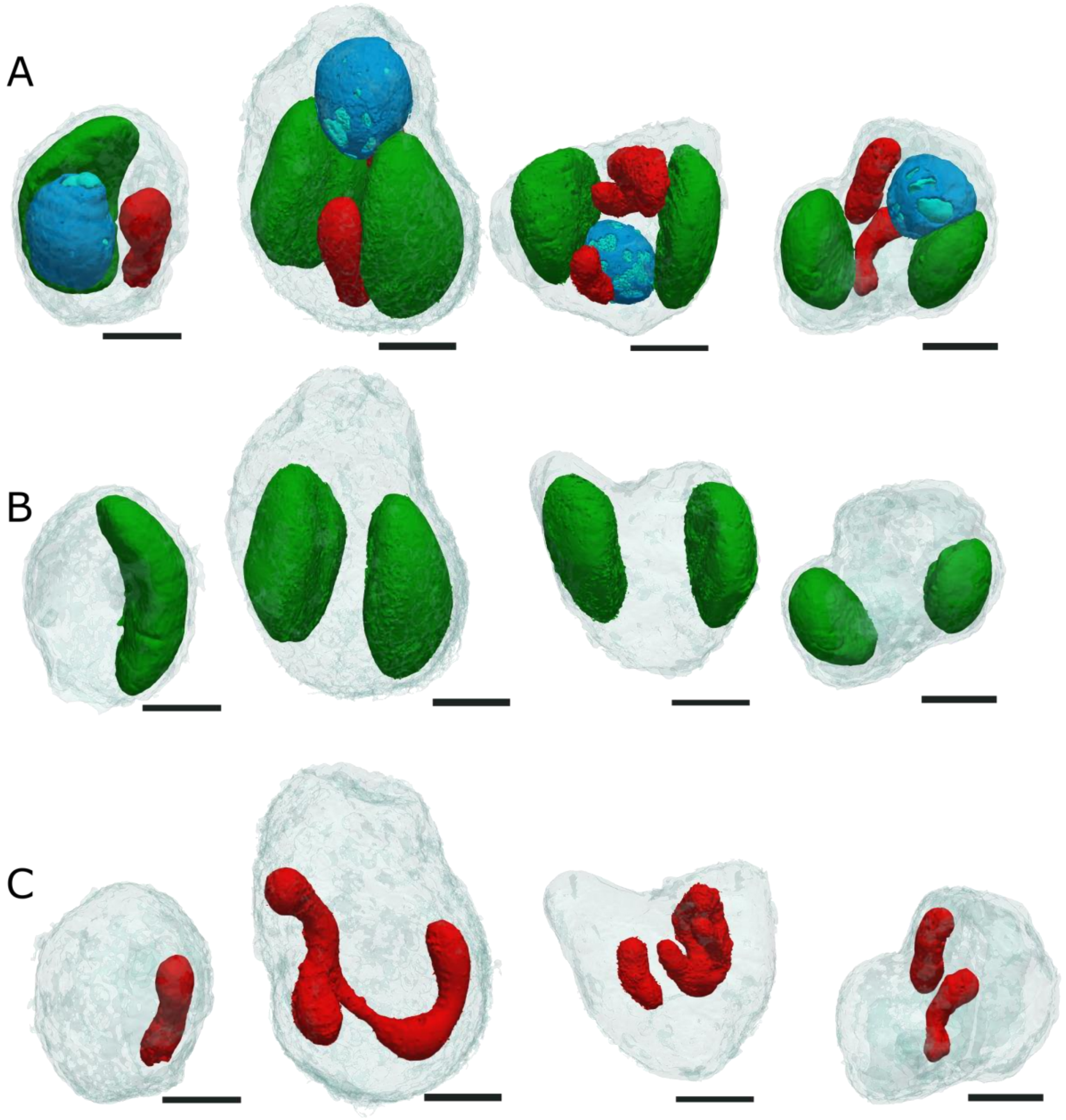

**Fig. S3. 3D reconstruction of five *Phaeocystis* cells in free-living unveiled by FIB-SEM** (the four cells are presented as columns)**.** 3D reconstruction shows the nucleus (blue, A), plastids (green, B), and the different topologies of the mitochondria (red; C). Scale bar: 1 µm.

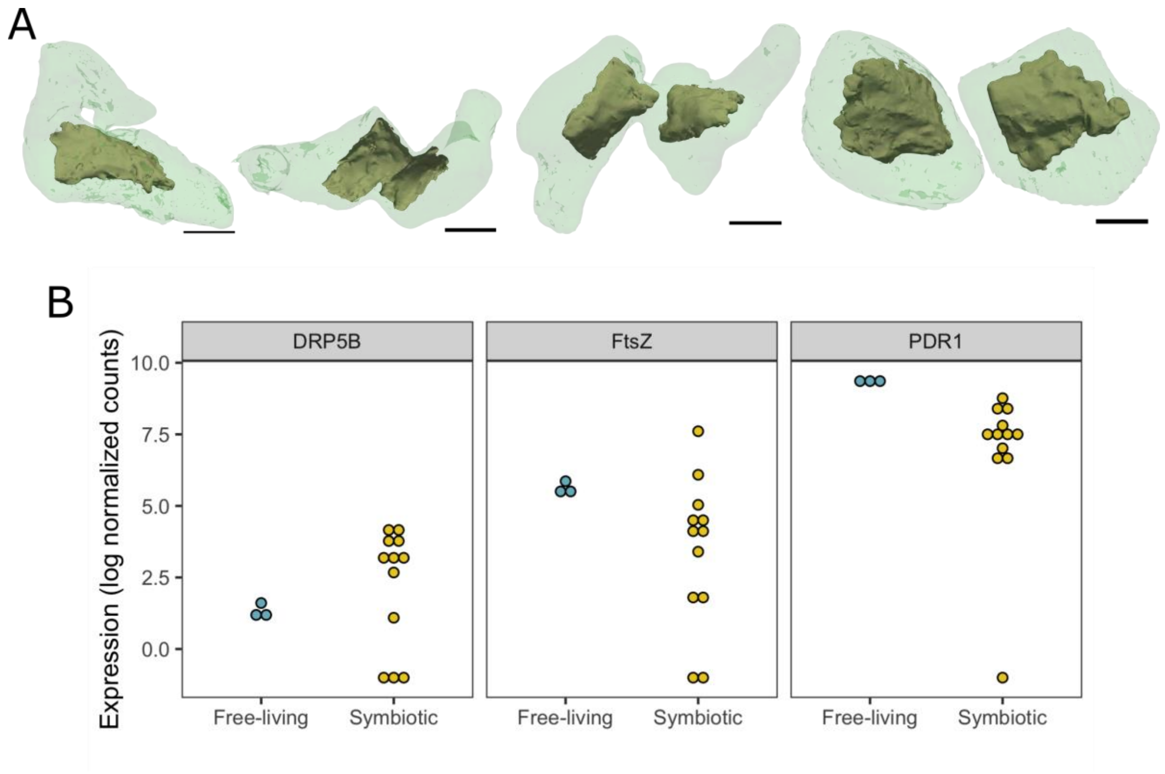

**Fig. S4. Division of plastids in *Phaeocystis* cells. (A)** Different steps of the plastid division as unveiled by FIB-SEM imaging in symbiotic *Phaeocystis* cells showing that the plastid (light green, transparency) enlarges and divides along with its pyrenoid (brown-green). Scale bar: 1 µm. **(B)** Expression levels of three genes (*DRP5B*, *FtsZ* and *PDR1*) involved in plastid division in *Phaeocystis* in the free-living and symbiotic stages. Points represent results from individual culture replicates for the free-living stage and individual host cells for the symbiotic stage.

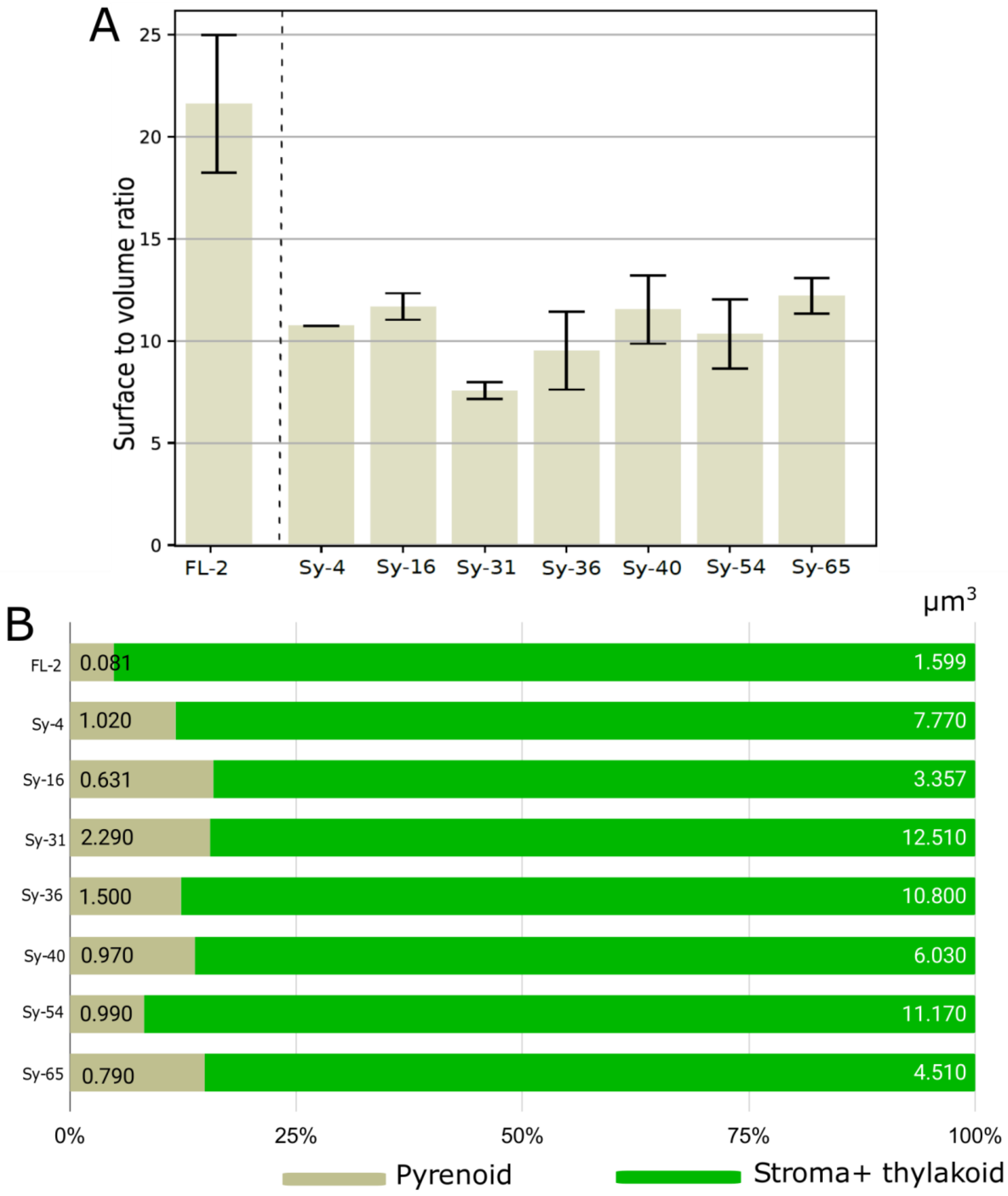

**Fig. S5. Morphometrics of the pyrenoid in free-living and symbiotic stage of *Phaeocystis*. (A)** Bar chart showing the average surface-volume ratio of the pyrenoid in free-living (n = 27) and in different symbiotic (Sy) cells with 4, 16, 31, 36, 40, 54, 65 plastids. Note that this surface-volume ratio is two fold higher in free-living cells compared to symbiotic ones. **(B)** Bar chart showing the volume occupancy of the pyrenoid in plastid, showing an increase in symbiosis (Sy-) compared to free-living *Phaeocystis* cells (FL-2).

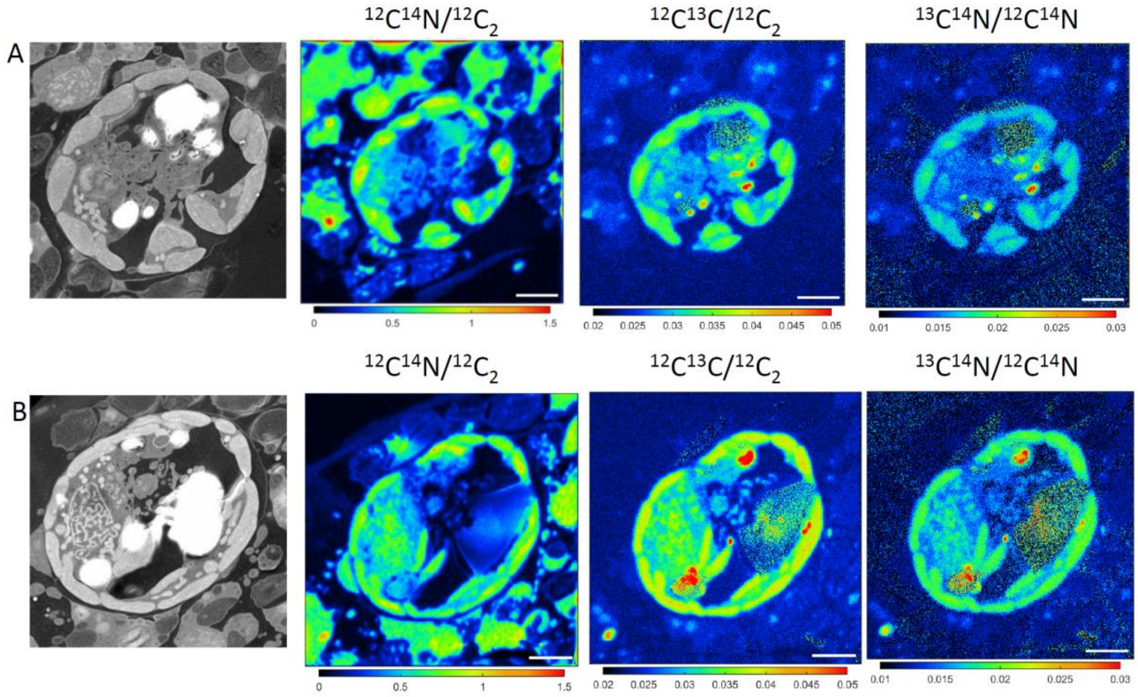

**Fig. S6. NanoSIMS (nanoscale Secondary Ion Mass Spectrometry) mapping of $^{13}C$ flux in symbiotic *Phaeocystis* cells (A and B).** Correlated SEM (Scanning Electron Microscopy) image (left) with nanoSIMS mapping of subcellular distribution of nitrogen ($^{12}C^{14}N/^{12}C_2$) and $^{13}C$ uptake (represented by the isotopic ratio $^{12}C^{13}C/^{12}C_2$) after 5 hours of incubation with $^{13}C$-labelled bicarbonate in filtered seawater.scale bar: 3 µm.

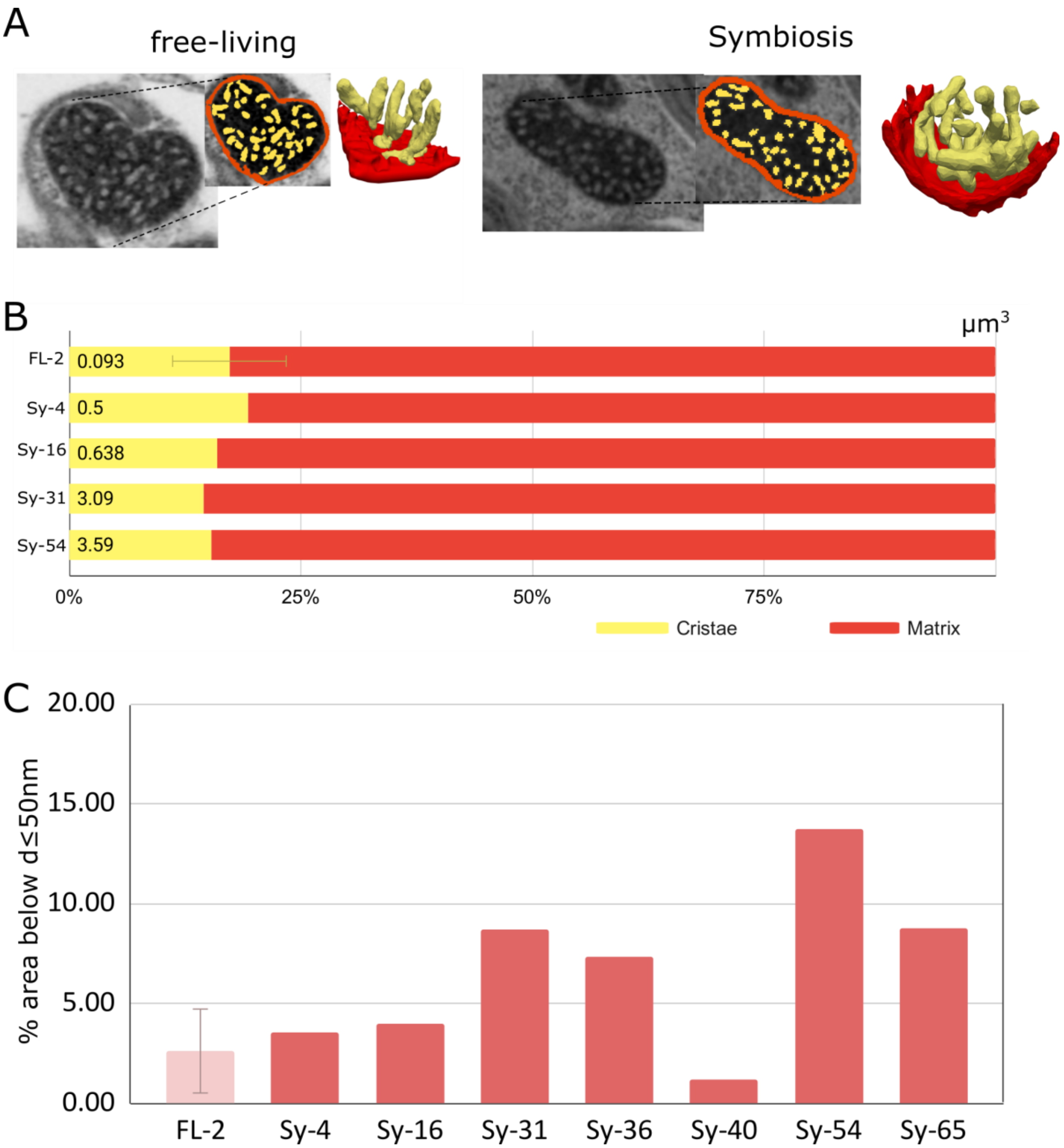

**Fig. S7. Morphometrics of the mitochondria and their cristae in free-living and symbiotic *Phaeocystis*, and interaction with the plastid. (A)** Morphology of the cristae in free-living and in symbiosis as unveiled by FIB-SEM imaging. **(B)** Bar chart showing the volume occupancy of the cristae in the mitochondria of the free-living (FL) stage (2 plastids, n = 14 cells) and four symbiotic cells (SY) with 4, 16, 31 and 54 plastids. **(C)** Bar chart showing the surface area (%) of the mitochondria in contact (< 50nm) with the plastid in free-living (2 plastids, n = 14), and in the symbiotic cells (Sy).

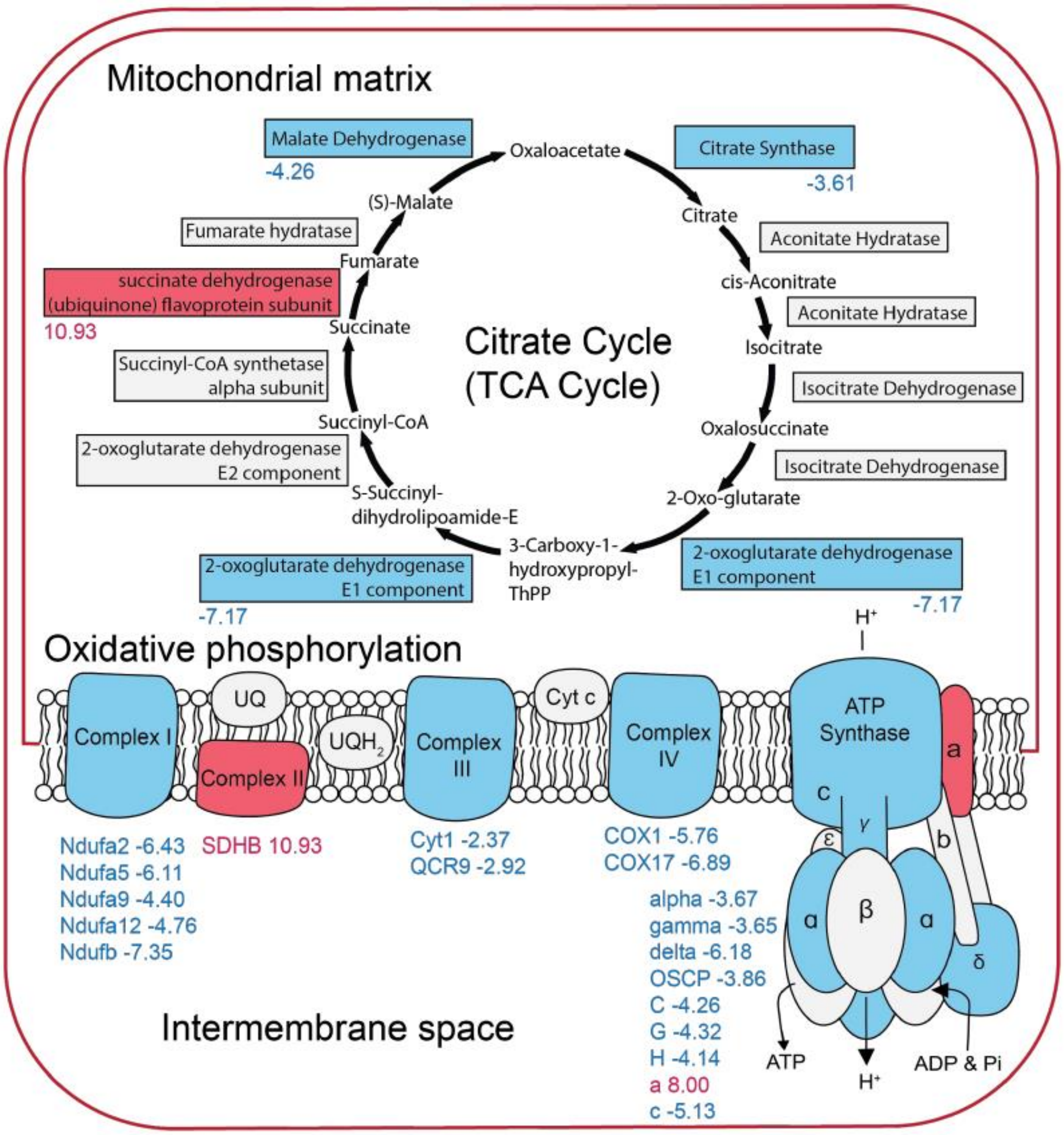

**Fig. S8. Genes involved in cellular respiration are primarily downregulated in symbiotic *Phaeocystis*.** Differential expression of genes in the Citrate Cycle (TCA Cycle) KEGG reference pathway (ko00020) and Oxidative Phosphorylation reference pathway (ko00190). Complexes and enzymes for which genes were significantly differentially expressed in symbiosis (log2FC > |1| and padj ≤ 0.05) are solid blue (downregulated; negative fold-change) or solid red (upregulated; positive fold-change). Genes for subunits in grey were not significantly differentially expressed in symbiosis. The log2FC for differentially expressed genes encoding enzymes or subunits are indicated below or within their representative shapes or blocks. The majority of genes in these respiratory pathways were downregulated in symbiotic cells.

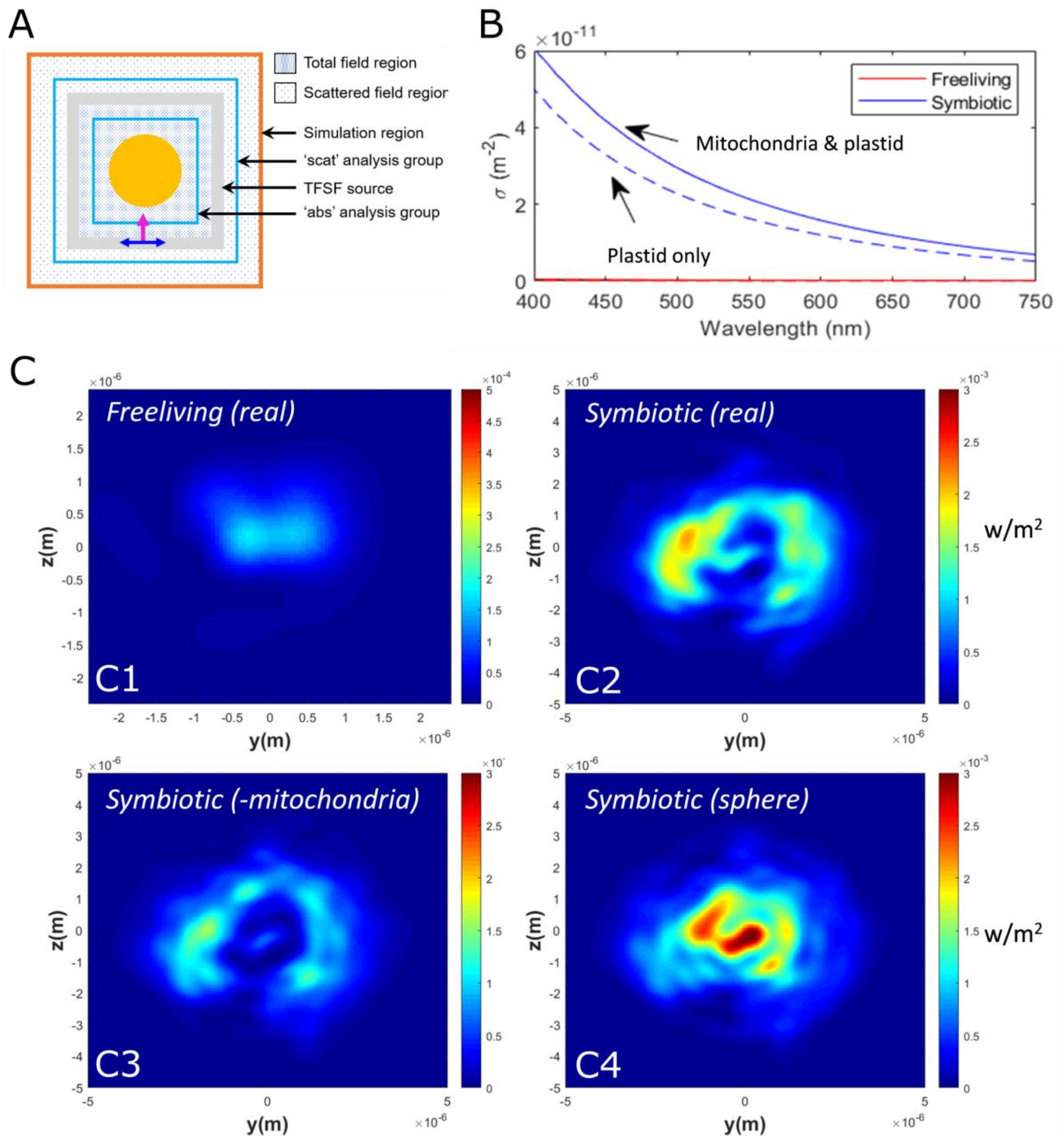

**Fig. S9. Photon propagation in free-living and symbiotic *Phaeocystis* using finite-difference-time-domain (FDTD) model. (A)** Computational set-up showing the Total-Field Scattered-Field (TFSF)-source, analysis groups and simulation regions (from https://support.lumerical.com/hc/en-us/articles/360042703433-Mie-scattering-FDTD-). **(B-C)** The scattering cross-section, σ **(B)** and the poynting vector **(C)** for free-living and symbiotic *Phaeocystis*. The real part of the poynting vector visualises the direction of power flow (w/m$^2$) over the *z,y* plane. The free-living cell (C1) shows a smaller power flow mainly in the forward direction. For the symbiotic case (C2) the power flow is higher and widely distributed. Removing the mitochondrial network reduces this effect (C3). Replacing the mitochondrial network with a sphere of the same volume with the same optical parameters (C4) shows that the morphology of the network strongly affects the light distribution.

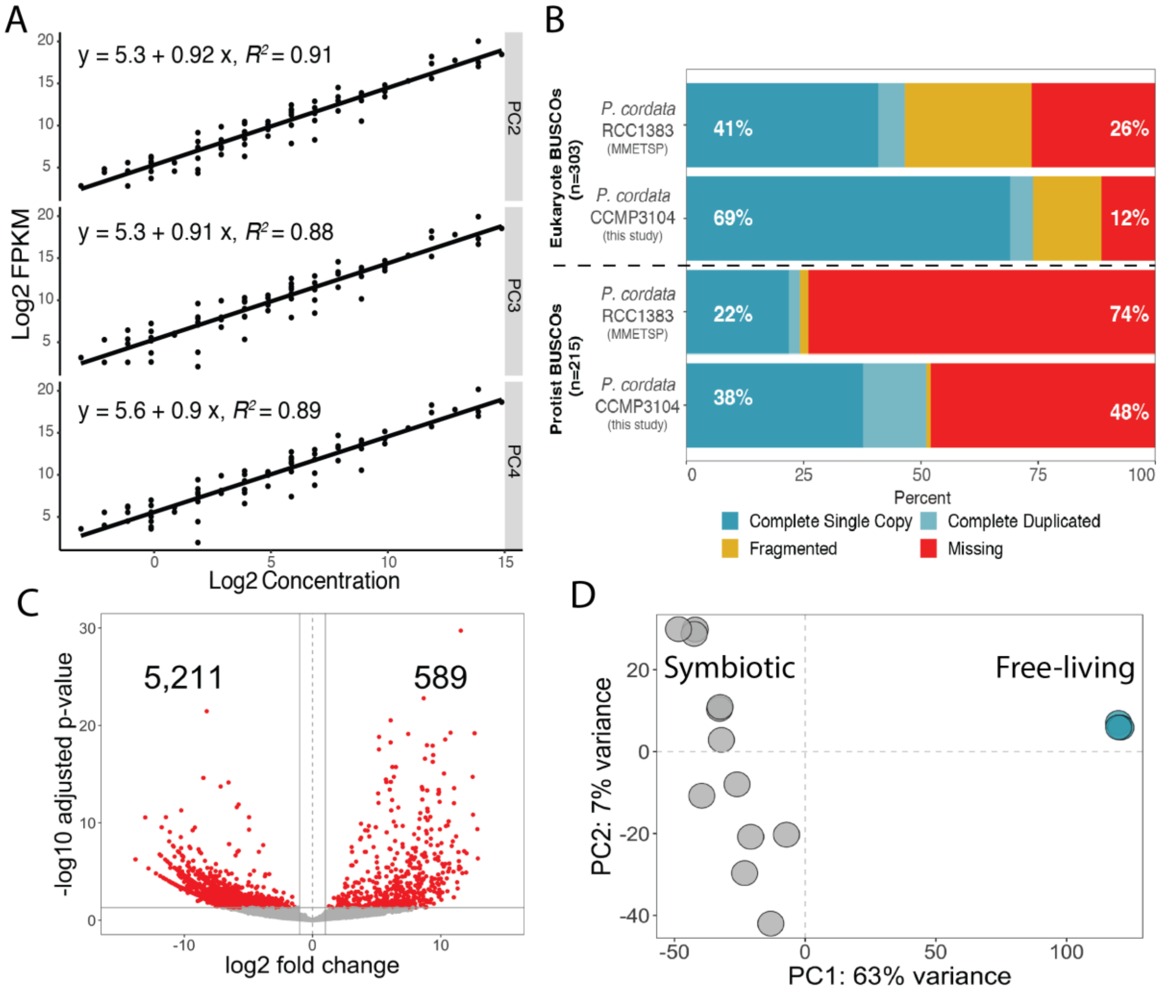

**Fig. S10. Transcriptomic results from free-living and symbiotic *Phaeocystis*. (A)** Linear regressions of External RNA Controls Consortium (ERCC) standard sequence initial concentrations and sequence counts. Quality-filtered sequences for each culture replicate (PC2–4) were mapped to ERCC standard sequences and read counts were determined with RSEM software. Fragments Per Kilobase per Million reads (FPKM) were log transformed and plotted against the log transformed initial ERCC concentrations. Regression lines, equations and $R^2$ values are included in each plot panel. **(B)** Percent of Eukaryotic and Protistan Benchmarking Universal Single-Copy Orthologs (BUSCOs) present in *Phaeocystis cordata* transcriptomes. The *P. cordata* RCC1383 transcriptome was previously sequenced as part of the Marine Microbial Eukaryote Transcriptome Sequencing Project (MMETSP). The *P. cordata* CCMP3104 transcriptome was sequenced for this study and is more complete than the previously assembled *P. cordata* transcriptome. **(C–D)** Results from differential gene expression testing between free-living and symbiotic *Phaeocystis cordata*. **(C)** Volcano plot displaying the relationship between log2 fold change in symbiotic samples and the False Discovery Rate (FDR) adjusted p-value calculated with the DESeq function. Genes with a log2 fold change greater than |1| and an adjusted p-value < 0.5 are considered significant and are colored red. The numbers of significantly downregulated genes (5,211) and upregulated genes (589) are indicated in the respective quadrants. **(D)** Principal Component Analysis (PCA) of log normalized read counts. Symbiotic samples are colored grey (n = 12) and free-living samples are blue (n = 3). The majority of variance is explained by sample-type.

**Table S1:** Results table from KEGG (Kyoto Encyclopedia of Genes and Genomes) pathway enrichment. Pathway is the KEGG pathway identifier; N is the total number of genes in the pathway that were annotated in the *P. cordata* transcriptome; DE is the number of genes in the pathway that were included in significantly up- or downregulated gene sets; p is the p-value reported by the kegga function in the edgeR package (only significant results are included); name is the KEGG name for the given pathway.

| | Pathway | N | DE | *p* | Name |
|---|---|---|---|---|---|
| Enriched among upregulated genes | ko03010 | 193 | 40 | 0 | Ribosome |
| | ko00195 | 28 | 9 | 0 | Photosynthesis |
| | ko00010 | 98 | 13 | 0 | Glycolysis / Gluconeogenesis |
| | ko00061 | 35 | 7 | 0.001 | Fatty acid biosynthesis |
| | ko00860 | 43 | 6 | 0.009 | Porphyrin and chlorophyll metabolism |
| | ko00053 | 7 | 2 | 0.032 | Ascorbate and aldarate metabolism |
| | ko00760 | 17 | 3 | 0.033 | Nicotinate and nicotinamide metabolism |
| | ko00190 | 75 | 7 | 0.038 | Oxidative phosphorylation |
| | ko05166 | 8 | 2 | 0.042 | Human T-cell leukemia virus 1 infection |
| | ko05162 | 1 | 1 | 0.042 | Measles |
| Enriched among downregulated genes | ko03030 | 42 | 17 | 0.001 | DNA replication |
| | ko00190 | 75 | 24 | 0.004 | Oxidative phosphorylation |
| | ko00480 | 34 | 13 | 0.006 | Glutathione metabolism |
| | ko00240 | 28 | 11 | 0.009 | Pyrimidine metabolism |
| | ko04146 | 33 | 12 | 0.013 | Peroxisome |
| | ko00051 | 22 | 9 | 0.013 | Fructose and mannose metabolism |
| | ko00531 | 45 | 15 | 0.014 | Glycosaminoglycan degradation |
| | ko00513 | 7 | 4 | 0.026 | Various types of N-glycan biosynthesis |
| | ko00052 | 45 | 14 | 0.031 | Galactose metabolism |
| | ko00790 | 18 | 7 | 0.037 | Folate biosynthesis |
| | ko04014 | 51 | 15 | 0.042 | Ras signaling pathway |
| | ko04742 | 8 | 4 | 0.045 | Taste transduction |
| | ko00534 | 8 | 4 | 0.045 | Glycosaminoglycan biosynthesis |
| | ko01523 | 15 | 6 | 0.046 | Antifolate resistance |
| | ko00780 | 5 | 3 | 0.048 | Biotin metabolism |

| | | | | | |
|---|---|---|---|---|---|
| | ko00361 | 5 | 3 | 0.048 | Chlorocyclohexane and chlorobenzene degradation |

**Table S2** *Phaeocystis cordata* CCMP3104 transcriptome assembly and annotation results

| | |
|---|---|
| **Number of contigs** | 32,621 |
| **Total number of base pairs** | 37,640,601 |
| **Minimum contig length** | 201 |
| **Maximum contig length** | 13,685 |
| **Median contig length** | 955 |
| **Mean contig length** | 1,153 |
| **N50** | 1,866 |
| **GC%** | 67.4% |
| **Contigs with Pfam annotation** | 13,548 (41.5%) |
| **Contigs with GO annotation** | 7,119 (21.8%) |
| **Contigs with KO annotation** | 7,719 (23.7%) |
| **Contigs with KEGG pathway annotation** | 4,618 (14.2%) |

**Movie S1 (separate file).** Stack of electron micrographs produced by FIB-SEM from the free-living stage of the microalga *Phaeocystis cordata* with 1 plastid.

**Movie S2 (separate file).** Stack of electron micrographs produced by FIB-SEM from the free-living stage of the microalga *Phaeocystis cordata* with 2 plastids.

**Movie S3 (separate file).** Stack of electron micrographs produced by FIB-SEM from the symbiotic stage of the microalga *Phaeocystis cordata* with 4 plastids.

**Movie S4 (separate file).** Stack of electron micrographs produced by FIB-SEM from the symbiotic stage of the microalga *Phaeocystis cordata* with 16 plastids.

**Movie S5 (separate file).** Stack of electron micrographs produced by FIB-SEM from the symbiotic stage of the microalga *Phaeocystis cordata* with 31 plastids.

**Movie S6 (separate file).** Stack of electron micrographs produced by FIB-SEM from the symbiotic stage of the microalga *Phaeocystis cordata* with 36 plastids.

**Movie S7 (separate file).** Stack of electron micrographs produced by FIB-SEM from the symbiotic stage of the microalga *Phaeocystis cordata* with 54 plastids.

**Movie S8 (separate file).** Stack of electron micrographs produced by FIB-SEM from the symbiotic stage of the microalga *Phaeocystis cordata* with 65 plastids.

**Dataset S1 (separate file).** Morphometrics of different organelles (volume and surface) of the microalga *Phaeocystis* in free-living and symbiotic stages calculated after FIB-SEM imaging. Proximity distance between plastid and mitochondria was also calculated.

**Dataset S2 (separate file).** Carbon uptake and $^{13}C$-enrichment of the symbiotic *Phaeocystis* in Acantharia and free-living *Phaeocystis* cells based on EA-IRMS analysis.
